# Ancestral circuits for vertebrate colour vision emerge at the first retinal synapse

**DOI:** 10.1101/2020.10.26.356089

**Authors:** Takeshi Yoshimatsu, Philipp Bartel, Cornelius Schröder, Filip K Janiak, Francois St-Pierre, Philipp Berens, Tom Baden

## Abstract

For colour vision, retinal circuits separate information about intensity and wavelength. This requires comparison of at least two spectrally distinct photoreceptors, as in the case of most mammals. However, many vertebrates use the full complement of four ‘ancestral’ cone-types (‘red’, ‘green’, ‘blue’, ‘UV’), and in those cases the nature and implementation of this computation remains poorly understood. Here, we establish the complete circuit architecture of outer retinal circuits underlying colour processing in larval zebrafish, which involves the full ancestral complement of four cone- and three horizontal cell types. Our findings reveal that the synaptic outputs of red- and green-cones efficiently rotate the encoding of natural daylight in a principal component analysis (PCA)-like manner to yield primary achromatic and spectrally-opponent axes, respectively. Together, these two cones capture 91.3% of the spectral variance in natural light. Next, blue-cones are tuned so as to capture most remaining variance when opposed to green-cones. Finally, UV-cones present a UV-achromatic axis for prey capture. We note that fruit flies – the only other tetrachromat species where comparable circuit-level information is available - use essentially the same strategy to extract spectral information from their relatively blue-shifted terrestrial visual world. Together, our results suggest that rotating colour space into primary achromatic and chromatic axes at the eye’s first synapse may be a fundamental principle of colour vision when using more than two spectrally well-separated photoreceptor types.

## INTRODUCTION

In visual scenes, information about wavelength is fundamentally entwined with information about intensity because the spectrum of natural light is highly correlated (*1*–*3*). Accordingly, wavelength information must be extracted by comparing the signals from at least two spectrally distinct photoreceptors, in a process generally referred to as “colour opponency” (*4*). To this end, most animal eyes use up to five spectral types of photoreceptors for daylight vision, with around four being the norm for vertebrates (reviewed in (*4*)). However, our knowledge of how the signals from four or more spectral types of photoreceptors are harnessed at a circuit level to extract this specific chromatic information remains limited.

Increasing the diversity of available spectral photoreceptors exponentially expands the diversity of theoretically detectable spectral contrasts. However, there is a law of diminishing returns: In natural scenes, some spectral contrasts are much more abundant than others. For efficient coding (*5*–*7*), animal visual systems should therefore prioritise the specific contrasts that are particularly prevalent in their natural visual world.

Here, we explored how zebrafish extract wavelength and intensity information from their natural visual world. Like many surface-dwelling fish, already their larvae use the ‘full’ ancient tetrachromatic cone-photoreceptor complement comprising red-, green-, blue- and UV-cones (*8*). Importantly, their retinal circuits can be non-invasively monitored and manipulated in the live animal (*9*) to provide insights into the computation of colour in the intact circuit.

We asked three questions: (i) What is the *in vivo* spectral tuning of zebrafish cone-outputs at the synapse, (ii) what is the circuit implementation, and (iii) how does this specific tuning support efficient sampling and decomposition of natural light?

Surprisingly, we found that two of the four cone types (green and blue) are strongly opponent, while the remaining two (red and UV) are essentially non-opponent, despite feeding into the same horizontal cell network. We go on to show how this spectral tuning is anatomically and functionally implemented at the circuit-level using horizontal cells. Further, comparison of the spectral tuning of the four cone-types to the spectral statistics of natural light showed that this specific cone-tuning arrangement allows zebrafish to effectively ‘solve’ a major fraction of the basic wavelength discrimination problem already at the first synapse of their visual system: Red-cones encode “colour-invariant” achromatic information, green-cones encode “brightness-invariant” spectral information, blue-cones provide a second chromatic axis that can be further optimised by possible opposition to green cones downstream, while UV-cones by themselves provide a secondary ‘UV-achromatic’ signal – presumably for visual prey capture (*10*). These findings also strongly imply that ancestral vertebrate circuits for colour vision are built upon the opponent signals from green- and blue-cones, which are lost in mammals including in humans (*4*).

Finally, zebrafish are not alone in using such an efficient strategy. By linking the spectral tuning of *Drosophila melanogaster* photoreceptors (*11*) with hyperspectral natural imaging data (*12*) we note that fruit flies use essentially the same strategy. However, their spectral tunings are systematically blue-shifted compared to those of zebrafish, presumably to directly acknowledge the relatively blue-shifted statistics of natural light in air (*13*). Taken together, our findings highlight a potentially general circuit-level mechanism of vision whereby incoming light is decomposed into “colour” and “greyscale” components at the earliest possible site.

## RESULTS

### Spectral tuning of zebrafish cones *in vivo*

To determine spectral tuning functions of the larval zebrafish’s four cone types (*8*) (red, green, blue, UV), we custom-built a hyperspectral full-field stimulator based on an earlier design (*14*) (Fig. S1a,b). A diffraction grating was used to reflect the light from 14 LEDs (peaks: 360 to 655 nm) into a collimated fibreoptic that was pointed at the live zebrafish’s eye mounted under a 2-photon (2P) microscope. To avoid spectral cross-talk with the 2P imaging system, we line-synchronised each LED’s activity with the scanner retrace (*15, 16*). Together, this arrangement permitted spectrally oversampling the much broader cone opsins (Fig. S1b,c) during *in vivo* 2P imaging in the eye. All stimuli were presented as wide-field flashes from dark.

### Green and blue cones, but not red and UV cones, display strong spectral opponency

We generated four cone-type specific SyGCaMP6f lines (Fig. 1a,b) (*17*) and measured the spectral tuning of each cone type at the level of their pre-synaptic terminals (pedicles), i.e. their output (Fig. 1c,d). Here, cones connect with other cones via gap junctions (*18*), with horizontal cells (HCs) which provide both feedback and feedforward inhibition (*19*), as well as with bipolar cells (BCs) which carry the photoreceptor signal to the feature extracting circuits of the inner retina (*20*). We did not study rods, as these are functionally immature in zebrafish larvae (*21, 22*).

**Figure 1.**
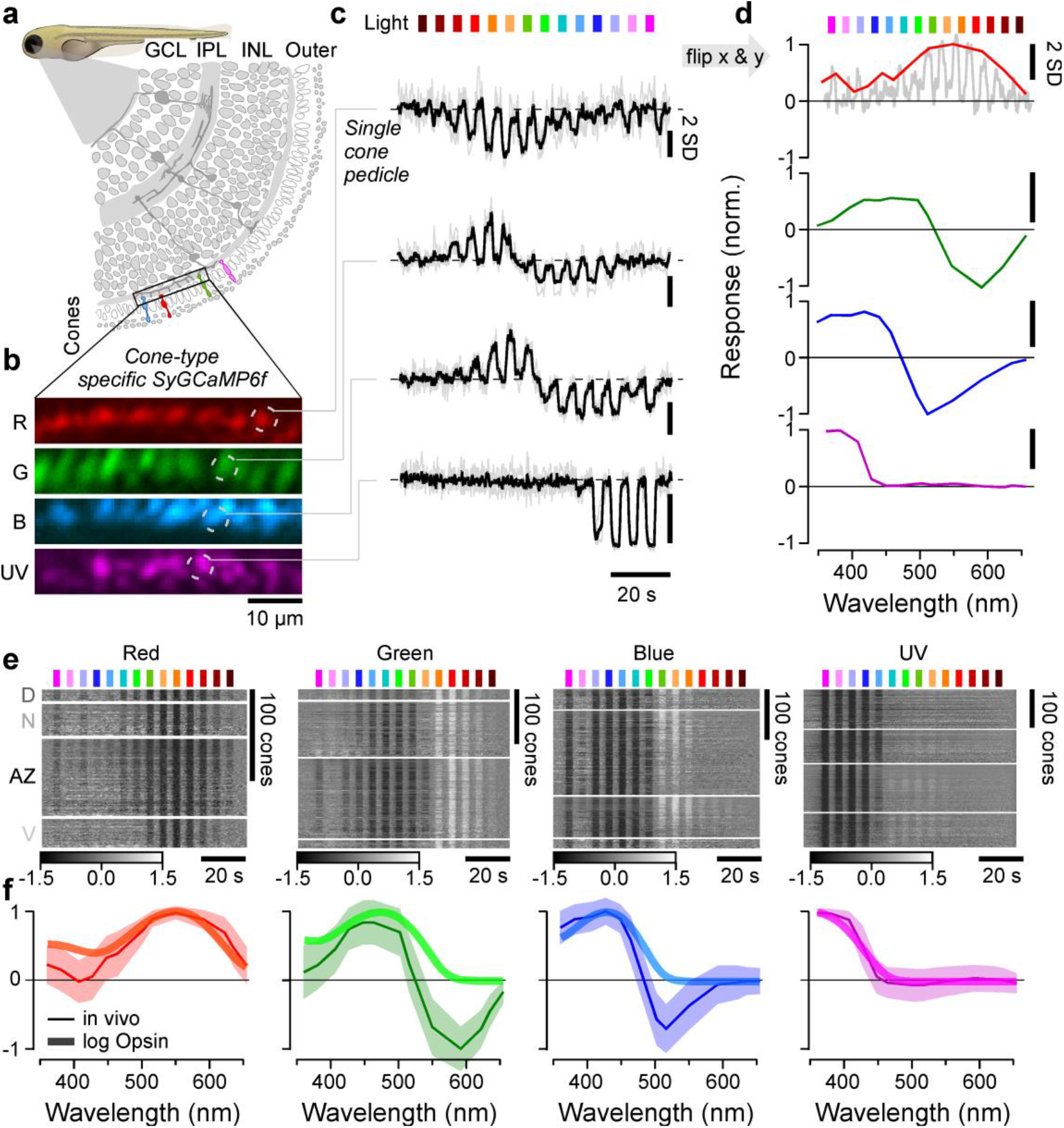
In vivo spectral tuning of larval zebrafish cones and HC block. **a**, Schematic of larva zebrafish retina, with position of cone-pedicles highlighted (adapted from (75)). **b,c**, example scans of the four spectral cones (b, Methods) with single pedicle response examples for each (c) to 3 s flashes of light from each of the 14 LEDs (see Fig. S1a-c). Shown are the means superimposed on individual repeats. **d**, Example spectral responses summarised from (c) – note that in this representation, both the X and y axes are flipped relative to the raw responses. **e**,**f**, Population responses of each cone types recorded in different parts of the eye (D, Dorsal; N, Nasal; AZ, Acute Zone; V, Ventral – see also Fig. S1g) (e) and population mean±95% confidence intervals with log-transformed respective opsin template superimposed (f, Methods). Heatmaps (e) are time-inverted to facilitate comparison to summary plots (f), greyscale bars are in z-scores.

From fluorescence traces, we extracted tuning functions (Methods), inverting both the x- and y-axes (Fig. 1d and inset). The inversions were done to display tuning functions from short- to long-wavelengths as is conventional, and to compensate for the fact that vertebrate photoreceptors hyperpolarise in response to light (*23*). We adhered to the time-inversion henceforth to facilitate comparison between raw data and summary plots (e.g. Fig. 1e). We systematically measured such tuning functions for n = 409, 394, 425, 431 individual red-, green-, blue- and UV-cones, respectively (n= 9, 11, 12, 7 fish). A total of n = 172, 288, 312, 410 recordings, respectively, passed a quality criterion (Methods, Fig. S1d-g) and were kept for further analysis.

Because the larval zebrafish eye is both structurally and functionally asymmetrical (*10, 13, 24*–*27*), we always sampled from four different regions of the eye’s sagittal plane: dorsal (D), nasal (N), ventral (V) and the *area temporalis* (acute zone, AZ (also known as “strike zone” (*13*))). With exceptions noted below (see also Discussion), we found that the spectral tuning of cones was approximately eye-position invariant (Fig. S1g). For further analysis we therefore averaged across cones irrespective their position in the eye (Fig. 1e,f).

On average, red- and UV-cones had approximately monophasic (non-opponent) output tuning functions that were largely in line with the tuning function of their respective log-transformed opsins (Methods). Such a log-transform is expected from the nature of signal transfer between outer segment phototransduction to synaptic calcium in the pedicle (*11, 28, 29*). Red-cones were broadly tuned and never exhibited opponency (Fig. 1f, left). In fact, some individual red-cones hyperpolarised in response to all tested wavelengths (Fig. 1e, left, cf. Fig. S1g). Nevertheless, on average red-cone sensitivity was weakly suppressed in the UV-range compared to the log-transformed opsin template (Discussion). In contrast, all UV-cones were narrowly tuned up to the short-wavelength cut-off imposed by the eye optics (∼350 nm, *unpublished observations*). Their tuning curve near perfectly matched the respective opsin template (Fig. 2f, right). UV-cones in the AZ and ventral retina moreover exhibited weak but significant opponency to mid-wavelengths (Fig. S1g, Discussion).

**Figure 2.**
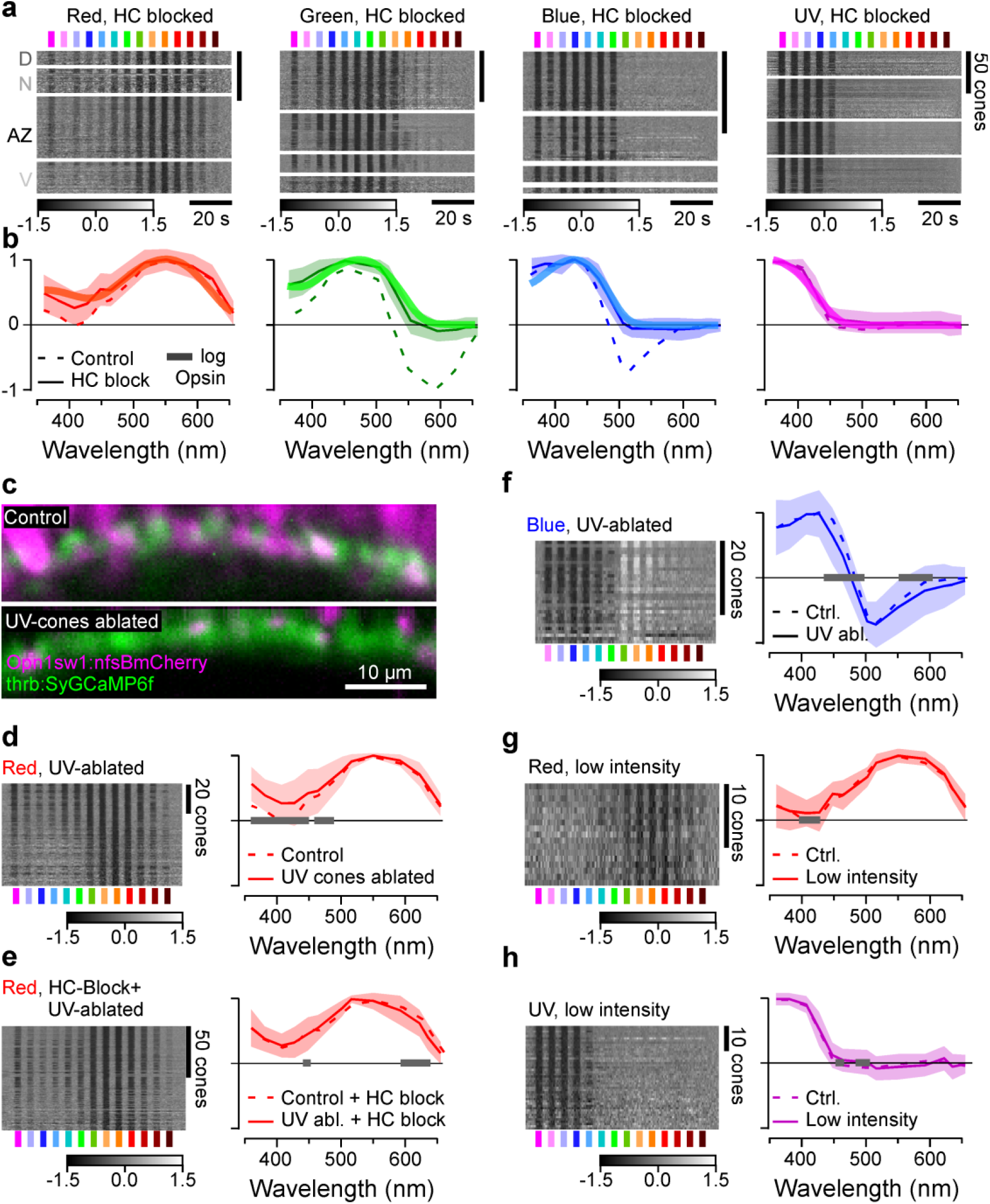
Opsin-like cone-responses in the absence of horizontal cells. **a**,**b**, Population responses of each cone type during pharmacological blockage of HCs (a, Methods) and population mean±95% confidence intervals with log-transformed respective opsin template superimposed (b, Methods). **c**, pharmaco-genetic UV-cone ablation in the background of red-cone GCaMP labelling before (top) and 24h after 2h treatment of metronidazole (10 mM) application (bottom, Methods). **d, e**, red-cone tunings after UV-cone ablation (n = 77) (d) and after additional pharmacological HC blockage (n = 103) (e). Shown are heatmaps (left) and means±SD (solid lines+shadings), and analogous data in the presence of UV-cones (dotted, from Figs. 1f, 2b). Note that the 361 nm LED was omitted in this experiment. **f**, as (d), but here recording from blue cones (n = 30). **g**,**h**, red- (n = 17) (g) and UV-cone tunings (n = 43) (h) at ∼9-fold reduced overall stimulus-light intensities (solid lines + shadings, Methods), compared to tunings at ‘standard’ light intensities (from Fig. 1f). Grey bars on the x-axis in (d-h) indicate significant differences based on the 99% confidence intervals of the fitted GAMs (Methods). Note that heatmaps (a,d-h) are time-inverted to facilitate comparison to summary plots (b, d-h). Grey-scale bars in z-scores.

Unlike red- and UV-cones, the *in vivo* output tuning functions of green- and blue-cones did not match their log-transformed opsin templates. Instead, these cones consistently exhibited strong spectral opponency to mid- and/or long-wavelength light (Fig. 1e,f, middle). Here, blue-cones had a highly consistent zero-crossing at 483±1 nm, while most green cones inverted at 523±1 nm (mean, 95% confidence intervals, Methods). Green-cones in the acute zone were slightly long-wavelength shifted with a zero-crossing at 533±1 nm (Fig. S1g, Discussion).

Together, to our knowledge this establishes the first direct and *in vivo* measurements of cone-pedicles’ spectral tuning functions in a vertebrate.

### Spectral tuning of zebrafish cones is fully accounted for by expressed opsins and horizontal cell feedback

The nature of phototransduction in cone-photoreceptors dictates that the absorption of photons leads to a drop in synaptic calcium. Accordingly, light-driven increases in synaptic calcium (Fig. 1f) must come from a sign-inverting connection from other cones, most likely via horizontal cells (HCs) (*30, 31*). We therefore decoupled HCs by pharmacologically blocking the glutamate output from cones using CNQX (Methods, Fig. 2a,b). This completely abolished all spectral opponency and increased the UV-response amplitude of red cones. As a result, now all four cone-tuning functions were fully accounted for by the respective log-transformed opsins (Fig. 2a,b, Fig. S2a). Our results further implied that possible heterotypical cone-cone gap junctions, if present, do not strongly contribute to spectral cone-tuning. In support, cone-tunings were essentially invariant to additional genetic ablation of UV-cones in the absence of HCs (Fig. 2c-f). Moreover, reducing overall stimulus brightness to probe for possible response saturation had no major effects on tuning functions (Fig. 2g,h). Taken together, our results strongly suggest that *in vivo*, the spectral tuning of all zebrafish cones is driven by the expressed opsin variant and shaped only by specific connections with HCs relaying feedforward signals from other cones. What are these HC connections?

### A connectome of the larval zebrafish outer retina

Light-microscopy studies in adult zebrafish have described at least three types of cone-HCs (H1-3), which contact R/G/B/(U), G/B/U and B/U cones, respectively (*30, 32*). However, for larval zebrafish HC-types and their connections to cones are not known except for H3 (*33*). To complete this gap in knowledge we used a connectomics approach based a combination of serial-section electron microscopy (Fig. 3) and confocal imaging (Fig. S3, Methods). In total, we reconstructed a 70 x 35 x 35 µm patch of larval outer retina in the acute zone, which comprised n = 140 cones n = 16 HCs (Fig. 3a-d). UV- and blue-cones were identified directly in the EM-volume based on their characteristic OPL-proximal mitochondrial pockets (UV, Fig. S3a) and somata (blue, Fig. S3b), respectively. This allowed initially sorting cones into three groups: UV, blue and red/green. Next, we traced each HC’s dendritic tree and identified their connections to cones belonging to each of these cone-groups (Fig. 3d-k, Fig. S3c-h). Relating each HC’s relative connectivity to UV-cones to their connections to red/green-cones allowed separating HCs into three groups (Fig. 3i, Fig. S3g), which were verified by clustering the HCs on all extracted features (Methods). These were dubbed H1, H2, and H3, based on their similarity to known adult HC types (*32, 34, 35*). The same classification was then further confirmed by confocal microscopy (Fig. S3d-h). Of these, H1 reliably contacted all red/green-cones within their dendritic field, while H2 systematically avoided approximately half of these cones. In line with confocal data (Fig. S3), this allowed disambiguating red-cones (contacted only by H1) from green-cones (contacted by both H1 and H2). With the exception of n = 14 of 66 red-green cones that could not be unequivocally allocated due to their location at the edge of the volume (yellow, counted as 0.5 red, 0.5 green in Fig. 3b,d), this completed cone-type identifications.

**Figure 3.**
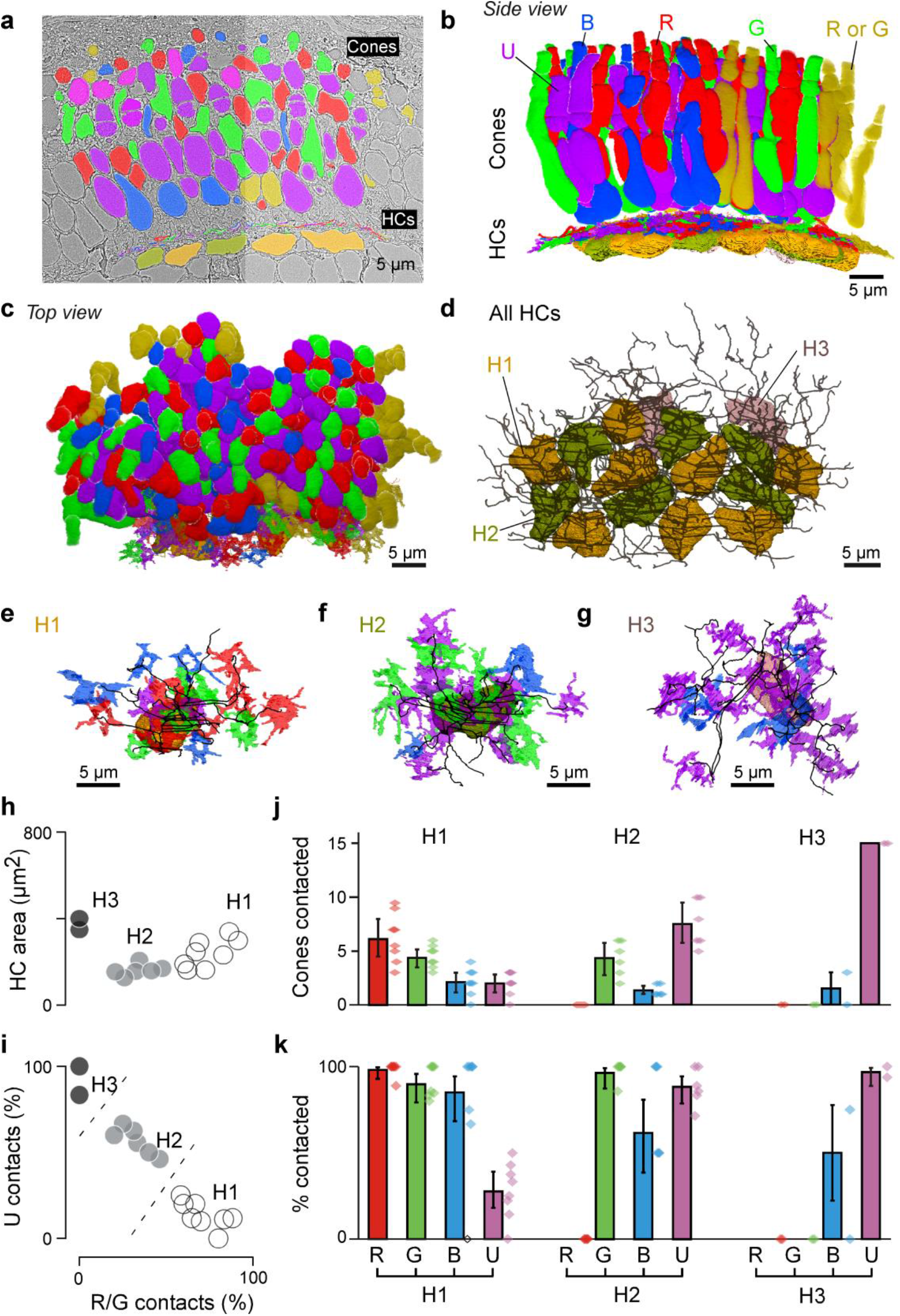
Connectomic reconstruction of outer retinal circuitry. **a**, Example vertical electron microscopy (EM) section through the outer retina, with cones and horizontal cells painted. Cones are colour coded by their spectral identity, with “yellow cones” indicating red- or green-cones at the section edge that could not be unequivocally attributed (Methods); HCs: H1, yellow/brown; H2, dark green, H3: light pink. **b-d**, Full volumetric reconstruction of all cones and skeletonised HCs in this patch of retina, shown from the side (b), top (c) and HC’s only (d). **e-g**, example individual HCs classified as H1 (e), H2 (f) and H3 (g) with connecting cone pedicles. **h-k**, Quantification of HC dendritic area (h, cf. Fig. S3g) and cone contacts (j-k) shown as absolute numbers with bootstrapped 95% CI (j) and percentage of cones in dendritic territory with binomial CI (i,k).

From here, we quantified each HC groups’ connections to the four cone types. This revealed that H1 contacted essentially all red-, green- and blue-cones within their dendritic fields, but imperfectly avoided UV-cones (Fig. 3j,k). In contrast, H2 by definition never contacted red-cones, but contacted all other cones including UV cones. Finally, H3 was strongly dominated by UV-cone contacts, with a small contribution from blue-cones. H3 never contacted red- or green-cones. Together, this largely confirmed that adult HC connectivity is already present in larvae, and moreover contributed cone-weighting information for the three HC types. We next asked how this specific HC-connectivity matrix underpins cone-spectral tunings.

### H1 horizontal cells likely underlie most spectral tuning

To explore how the three HC-types contribute to spectral cones-tunings, we first set up a series of functional circuit models for all possible combinations of HCs (Methods). These linear models included the established connectivity structure (Fig. 3k) and were driven by the cone tunings in the absence of HCs (Fig. 2a), with the goal of explaining cone-tunings in the presence of HCs (Fig. 1f). We computed posteriors for the model parameters using likelihood-free inference (*36*) based on the cones’ tunings, and we assumed sign-preserving connections from cones to HCs but sign-inverting connections from HCs to cones.

The model recapitulated well the *in-vivo* tuning functions of all cones when simultaneously drawing on all three HC types. However, almost the same fit quality was achieved when using H1 alone (Fig. 4a-d, cf. Fig. S4a-c), while H2 mainly fine-tuned the blue- and UV-cones and H3 had negligible impact on any cone-tunings (Fig. S4a). In fact, any model that included H1 outperformed any model that excluded H1 (Fig. S4a-c). H1, where present, also consistently provided the strongest feedback amongst HCs (Fig. 4d, Fig. S4c). Together, modelling therefore suggests that H1-like HCs are the main circuit element underlying the *in-vivo* spectral tuning of zebrafish cones. Moreover, the inferred relative cone-type weighting for H1 approximated their anatomical connectivity established by EM (Fig. 4j), with the exception of green-cones which had stronger-than-expected weights (Fig. 4d) – possibly uncovering an increased synaptic gain at this site.

**Figure 4.**
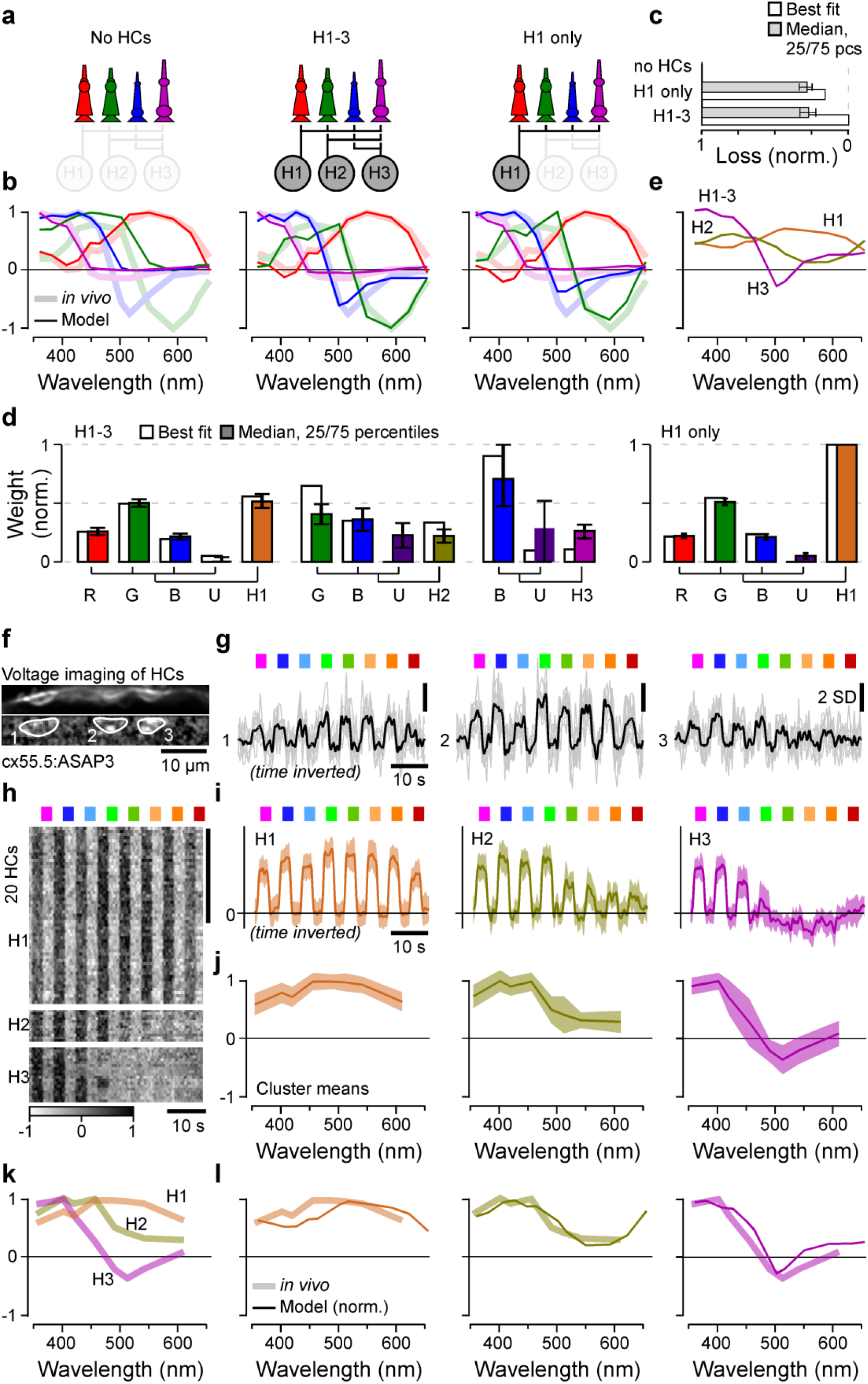
Spectral tuning of cones by horizontal cells. **a-e**, linear model of spectral tuning in an outer retinal network comprised of 4 cone- and 3 HC-types, with maximum connectivity matrix defined as in Fig. 3k (Methods). Cone tunings are initiated based on in vivo data during HC block (Fig. 2b). Different HC combinations include (a, from left): no HCs, all HCs and H1 only. In each case, the model computes resultant cone-tunings (solid lines) superimposed on in-vivo data in the absence of HC block (shadings, from Fig. 1f) (b), reconstruction quality (c) as loss relative to the peak performance for the full H1-3 model (loss = 0) and in the absence of HCs (loss = 1) and normalised weights such that cones contributing to a given HC, and HCs contributing to the full model, each add up to 1 (d).In addition, resultant HC tunings are shown for the full H1-3 model (e). **f-j**, in vivo voltage imaging of HC somata’s spectral tuning (Methods). f,g example scan (f, average image (top) and local response correlation (76) and ROIs (bottom)) and responses (g, mean superimposed on individual repeats). **h-j**, results of clustering of mean responses from n = 86 ROIs (h, n = 15 fish) with cluster means (i) and extracted tuning functions (j, means±SD). **k**,**l**, mean tunings of in vivo HC clusters (k, from j), and superposition of each measured modelled (solid lines, from e) and measured (shading, from k) HCs. Note that raw- (g) and averaged (i) HC-responses as well as the summary heatmap (h) are time-inverted to facilitate comparison with summary plots (j-l). Greyscale bar in (h) in z-scores.

Next, we sought to verify the model by experimentally measuring the spectral tunings of HCs and comparing these to the predicted HC tunings from the full model (Fig. 4e). For this, we used *in vivo* 2P voltage imaging of HCs somata using ASAP3 (*37*) (Fig. 4f-l) (Methods). In total, recordings from n = 86 HCs that passed a quality criterion (Methods) were sorted into three clusters (Methods). The largest cluster exhibited a spectrally broad, monophasic response that closely matched the model’s prediction for H1 (Fig. 4l, see also (*30, 34*)). Next, short-wavelength biased clusters 2 and 3 closely matched the model’s prediction for H2 and H3, respectively (*30, 34*).

### Efficient encoding of achromatic and chromatic contrasts in natural light

To explore how the specific *in vivo* cone tuning functions may support zebrafish vision in nature, we next computed the distribution of achromatic and chromatic content of light in their natural habitat. For this, we used a total of n = 30 underwater hyperspectral images (1,000 pixels each: 30,000 spectra) (*12, 13*) (Fig. 5a-c). Using one example scan for illustration (Fig. 5a), we first computed each cone’s view of the world in the absence of outer retinal feedback by taking the dot product of each log-transformed opsin spectrum with each pixel spectrum (Fig. 5d-f).

**Figure 5.**
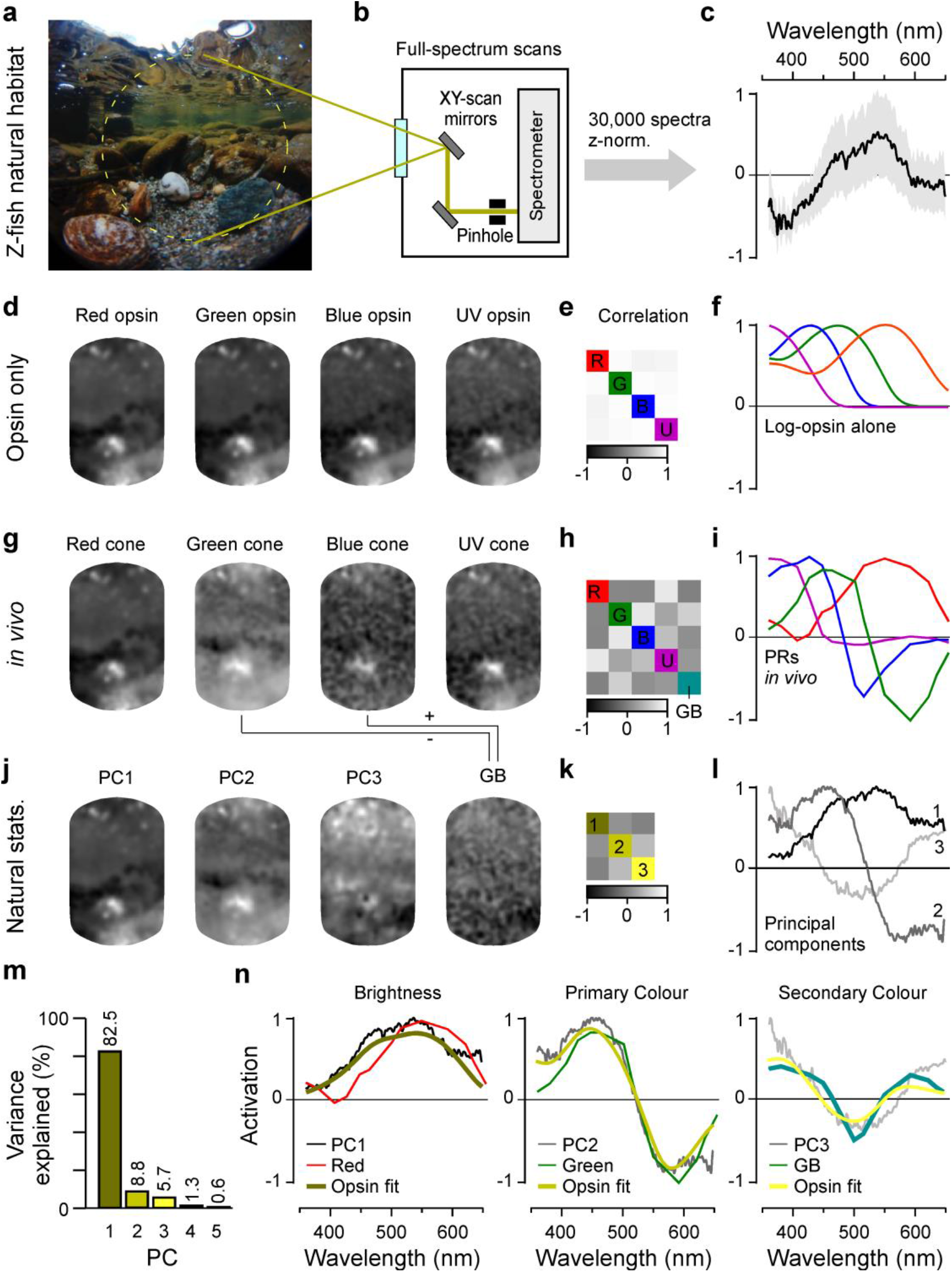
In vivo cone tunings efficiently represent statistics of natural light. **a-c**, Hyperspectral data acquisition from zebrafish natural visual world. A 60° window around the visual horizon of an example scene recorded in the zebrafish natural habitat (a) was sampled at 1,000 equi-spaced points with a custom-built spectrometer-based scanner (12) (b) to yield 1,000 individual spectral readings from that scene. (c) summarises the pooled and z-normalised data from n = 30 scenes (30,000 spectra) with mean±SD (data from (13)). **d-l**, reconstructions and analysis of the example scene as see through different spectral filters: (d-f) log-opsin spectra, (g-i) cone in vivo tunings and (j-l) based on first three principal components (PCs) that emerge from the hyperspectral data shown in (c). From left to right: (d,g,j) example scene (from a) reconstructed based on opsin-/in vivo-/PC-tunings as indicated, (e,h,k) correlation matrices between these respective reconstructions and (f,i,l) the actual tunings/PCs. A 5^th^ element “GB” (for “green/blue”) is computed for in vivo tunings as contrast between green- and blue-cone tunings (cf. Fig. S5). **m**, % variance explained by the first five principal components (l). **n**, Superposition of cone in-vivo tunings (coloured lines), PCs, and a linear R/G/B/U log-opsin fit to the respective PC (yellows, Methods). The latter fit can be seen as the biologically plausible optimum match to a given PC that can be achieved in a linear regime.

In this configuration, the intensity-normalised representations of the scene by each of the four cones were extremely similar as expected from high spectral correlations in natural light (Fig. 5d). In contrast, when the same scene was computed for the intact outer retinal network by taking the *in vivo* cone-tuning functions (from Fig. 1f), the different cones instead delivered much more distinct images (Fig. 5g-i).

Next, to determine the spectral axes that optimally captured the variance of natural light in the zebrafish’s natural underwater world (Discussion), we used principal component analysis (PCA) across the spectra of all n = 30,000 pixels in the data set (Fig. 5c, j-l). Due to the strong spectral correlations in natural light, the first component (PC1) captured the achromatic (“black and white”) image content, while subsequent components (PC2, PC3 etc.) captured the major chromatic (“colour”) axes in decreasing order of importance (*1, 5*). Together, PCs 1-3 accounted for 97% of the natural spectral variance (Fig. 5m). We computed what the example scene would look like if sampled by detectors that were directly based on the first three principal components. We found that scenes processed by PC1 and PC2 (Fig. 5j) were highly reminiscent of the scenes sampled by *in vivo* red- and green cones, respectively (Fig. 5g). Next, PC3 was not obviously captured by either of the remaining blue- or UV-cones in isolation, however it did approximately resemble the scene when reconstructed by a green/blue-cone opponent axis (“GB”, turquoise, Discussion). In fact, PC3 could be approximated by a variety of cone-combinations, however all best-matches (ρ=0.97, Methods) required opposing green- and blue-cones (Fig. S5).

Direct superposition of these cone-output spectra with the respective principal components further illustrated their striking match (Fig. 5n). These cone-spectra were also well matched by a direct fit to the principal components when using the four cones’ opsin-templates as inputs (Fig. 5n, yellows, Methods). Here, our rationale was that these opsin-fits present a biologically plausible optimum for mimicking the principal components.

To quantitatively explore this match and its consequences for the encoding of natural light, we next computed how each of the 30,000 individual collected spectra would activate red- and green-cones as well as the GB-axis. We then plotted these activations against the respective loadings of PC1-3 for these spectra (Fig. 6a). In each case, we also computed the same metric for the best log-opsin fits to the PCs. This confirmed the excellent performance of the system for separating achromatic from chromatic information under natural light. Red-cone activation correlated almost perfectly (mean ρ>0.99, 2.5/97.5 percentiles 0.99/>0.99) with spectral loadings against PC1 (Fig. 6a, top left, cf. Fig. 6b, top left), but was uncorrelated with either PC2 (ρ=-0.16, −0.89/0.88) or PC3 (ρ=0.29, − 0.34/0.91) (Figs. 6a,b, middle and bottom left). Moreover, red-cone performance was near-indistinguishable from that of the opsin fit against PC1 (ρ>0.99, >0.99/>0.99), which was used as a biologically plausible benchmark of optimality (Figs. 6a,b, second column). Accordingly, and despite the minor differences in short-wavelength activation of the red-cone action spectrum compared to PC1 and its opsin fit (Fig. 5n, left, Discussion), red-cones encoded natural achromatic contrast (i.e. “brightness”, PC1) with negligible contamination of chromatic information (i.e. PCs 2,3). In contrast, activation of green-cones was highly correlated with PC2 (ρ=0.99, 0.98/>0.99), but uncorrelated with either PC1 (ρ=-0.15; − 0.88/0.88) or PC3 (ρ=0.14, −0.66/0.81, Figs. 6a,b columns 3). Again, their performance was near-indistinguishable from that of the respective opsin fit (Figs. 6a,b, columns 4). Accordingly, green-cone activation carried no information about brightness, but instead encoded an efficient primary chromatic signal.

**Figure 6.**
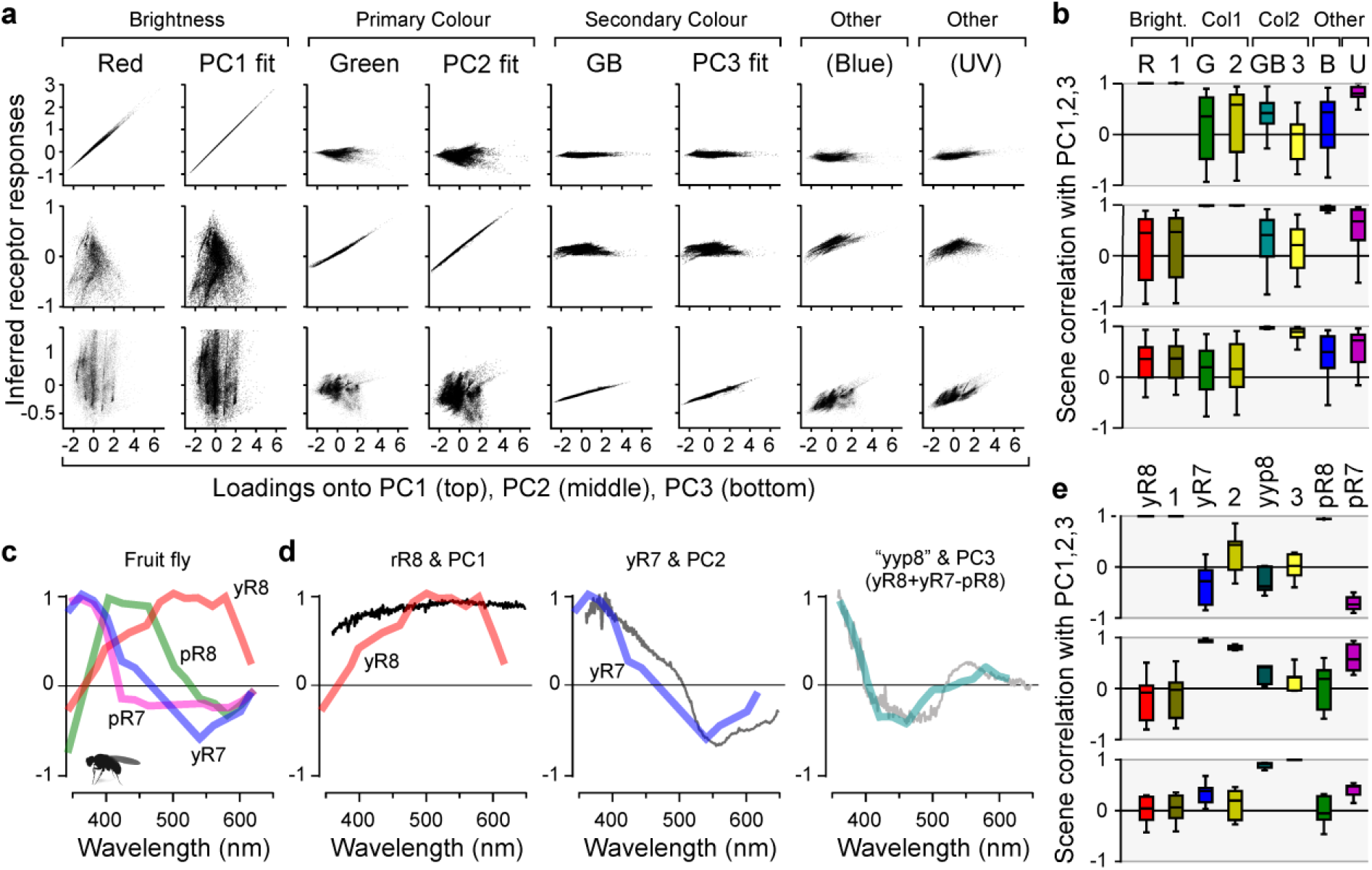
Encoding of natural achromatic and chromatic contrast. **a**, Computed “responses” of in vivo cones, the GB-axis, and each respective log-opsin PC-fit (all from Fig. 5i,n) to each of the n = 30,000 individual natural spectra, plotted against (each spectrum’s loadings onto PC1 (top row), PC2 (middle row) and PC3 (bottom row), as indicated. “Responses” plotted on y-axes, PC-loadings on x-axis. In general, a column that shows a near-perfect correlation in one row, but no correlation in both other rows (e.g. column 1) can be seen as a tuning function that efficiently captures the respective PC (e.g. column 1 shows that red-cones efficiently represent PC1 but not PC2 or PC3). **b**, Corresponding summary statistics from (a), based on scene-wise Spearman-correlations. **c**, Spectral tuning functions of Drosophila R7/8 photoreceptors as measured in vivo at their synaptic output (data from (11)). **d**, comparison of Drosophila tuning functions with the first three PCs that emerge from terrestrial natural scenes (data from (12)). Here, PC3 is matched with a “yyp8” axis as indicated (cf. Fig. S6d-f). **e**, Summary stats of Drosophila photoreceptor “responses” to each of the n = 4,000 individual terrestrial natural spectra plotted against their respective PC loadings.

Next, both activation of the GB-opponent axis and of the corresponding opsin fit correlated strongly with PC3 (ρ=0.95, 0.80/0.99; ρ=0.79, 0.18/0.99, respectively), but not with PC1 (ρ=0.38, −0.25/0.92; ρ=-0.08, − 0.76/0.52) or PC2 (ρ=0.31, −0.62/0.91, ρ=-0.15, −0.80/0.57, Figs. 6a,b, columns 5,6). Accordingly, contrasting the signals of blue- and green-cones offers the theoretical possibility to build an efficient secondary chromatic signal in downstream circuits (Discussion). Notably, blue-cones in isolation correlated mainly with PC2 (ρ=0.94, 0.85/0.99) rather than PC1 (ρ=0.18, −0.74/0.91) or PC3 (ρ=0.43, −0.33/0.90) (Figs. 6a,b, columns 7), suggesting that they could potentially serve to provide an alternative route to encoding primary chromatic information.

Finally, UV-cones mainly correlated with PC1 (ρ=0.80, 0.51/0.99), suggesting that this ultra-short-wavelength channel may serve to provide a secondary achromatic signal (Figs. 6a,b, columns 8). However, its performance in doing so was substantially inferior to that of red-cones, suggesting that its primary function is not the encoding of achromatic brightness *per se*, but rather to specifically detect short-wavelength signals. Here, their weak but significant opponency to spectrally intermediate signals may serve to accentuate contrast against an otherwise “grey” background (Discussion).

Taken together, it appears that larval zebrafish effectively ‘rotate’ colour space already at their visual system’s first synapse signal along an achromatic axis (red-cones) and a primary chromatic axis (green-cones), with the added possibility to build an efficient secondary chromatic axis by opposing green- and blue-cones downstream. Together, this system captures at least 91.3% of spectral variance in natural scenes when using red- and green-cones alone, and potentially up to 97% if including green-blue opponency. Elegantly, it also leaves UV-cones to serve independent visual functions, such as prey capture of UV-bright microorganisms (*10*) (Discussion).

### A comparison to spectral processing in fruit flies

A conceptually similar decomposition of natural light may also be used in *Drosophila melanogaster* (Fig. 6c-e, Fig. S6), the only other tetrachromatic species where *in vivo* spectral tuning functions of photoreceptor outputs are available (*11*). In these flies, R1-6 photoreceptors express a mid-wavelength sensitive opsin and are generally considered an achromatic channel, while R7/8-type photoreceptors are associated with colour vision (*38*). We therefore compared spectral tuning curves of the four varieties (yR8, yR7, pR8, pR7) of *Drosophila* R7/8-type photoreceptors (Fig. 6c, taken from (*11*)) with the principal components that emerged from natural spectra of n = 4 daytime field and forest scenes (*12*), each comprising 1,000 individual spectra as before (Fig. 6d,e, Fig. S6a-g, Discussion).

Like for zebrafish, this showed that their spectral tuning curves were well approximated by the first three terrestrial PCs: PC1 and yR8 (ρ>0.99, 0.99/>0.99), PC2 and yR7 (ρ=0.93, 0.91/0.98) and finally PC3 by opposing jointly opposing both yR8 and yR7 against pR8 (for simplicity: “yyp8”, ρ=0.72, 0.60/0.84, Fig. 6d,e, cf. Fig. S6d-g). Compared to zebrafish, the spectral matches between photoreceptor action spectra and natural PCs were however slightly worse, which may in part be linked to the use of a smaller natural imagery dataset, and to the comparatively lower spectral resolution information currently available in flies. Nevertheless, this general match was made possible by the fact that, in line with the relatively increased predominance of short-wavelength light above the water (Fig. S6a), all terrestrial principal components (Fig. S6b) and corresponding action spectra (Fig. S6g) were blue-shifted relative to those of aquatic environments and of zebrafish, respectively.

Together, this suggests that ‘rotating’ colour space into primary achromatic and chromatic axes (i.e. PC1-2) as early as possible, while leaving the ultra-short wavelength system largely isolated, may be a fundamental principle of colour vision when using more than two spectrally well-separated photoreceptor types, in a striking example of convergent evolution (Discussion).

## DISCUSSION

Our physiological recordings from cones (Figs. 1,2) and horizontal cells (Fig. 4f-l), linked to synaptic level EM-reconstructions (Fig. 3) and computational modelling (Fig. 4a-e) provide a comprehensive *in vivo* account of spectral processing for an efficient decomposition of natural light (Figs. 5,6) at the visual system’s first synapse in a tetrachromatic vertebrate.

### Linking retinal colour opponency to the principal components of natural light spectra

Using PCA of light spectra for understanding the encoding of natural scenes by animal visual systems has a long tradition, for example in information-theoretic considerations by Buchsbaum and Gottschalk in 1983 (*1*). This seminal work described how the three primaries of the human eye (long- mid- and short-wavelength sensitive: L/”red”, M/”green”, S/”blue”, respectively) can be efficiently combined to derive one achromatic and two chromatic axes with none, one and two zero crossings, respectively. These theoretically optimal channels corresponded well to psychophysically determined opponent mechanisms in human vision, and were later shown to capture much of the spectral variance in natural light (*3*). However, in contrast to zebrafish, the circuit mechanisms that enable this striking link between the human primaries and perception involve multiple levels of computation across both the retina and the brain remain incompletely understood: First, many retinal ganglion cells (RGCs) and their central targets, including in visual cortices, are mid/long-wavelength-biased and non-opponent, and encode achromatic contrasts (*39*). Second, inherited from probably non-selective retinal wiring, midget circuits carry “red-yellow” or “green-yellow” spectral information that is thought to be decoded into a primary “red-green” colour-opponent axis in the central brain by mechanisms that remain largely unsolved. Third, at least three types of “blue-yellow” RGCs contrast the signals from blue-cones against the sum of red- and green-cones. This RGC opponency is mainly achieved at the level of RGC dendrites, by contrasting the signals of approximately non-opponent inner retinal neurons (*40*).

In addition, primate blue-cones themselves are yellow-blue opponent due to feed-forward inputs of red-/green-cone inputs via HCs (*41*) – reminiscent of the strategies employed by zebrafish cones. However, primate blue- and red-/green-cones are homologous to zebrafish UV- and red-cones, respectively (*42, 43*), and the HC underlying is H2 (*40*). Accordingly, spectral opponency in primate blue-cones is presumably linked to the weak but significant mid-wavelength opponency of zebrafish UV-cones, rather than the much stronger opponency of zebrafish green- or and blue-cones (Fig. 1f).

Beyond primates, comparative circuit knowledge of vertebrate retinas for spectral processing is sparse and mainly restricted to dichromatic mammals (*4*). Amongst tetrachromats that retain ancestral green- and blue-cones, measurements of spectral responses in adult HCs of diverse species of fish (*30, 34, 44*) are in good agreement with our *in vivo* HC data in larval zebrafish. Moreover, zebrafish inner retinal neurons (*8, 13, 24, 45*) display both non-opponent as well as a wide diversity of opponent responses that generally prioritise simple short-vs.-long wavelength computations over more complex combinations, broadly in agreement with predictions from theory (*1*). However, in the absence of systematic and spectrally resolved sensitivity measurements of zebrafish inner retinal neurons, it has not been possible to explicitly link their properties to the variance in natural visual light. In addition, direct *in vivo* spectral measurements of zebrafish cone-photoreceptor outputs have remained outstanding.

Amongst invertebrates, *Drosophila melanogaster* stands out as the only tetrachromatic species where spectrally resolved photoreceptor output tuning functions are available (*11*). As discussed, these reveal a conceptual match to those of zebrafish, even down to circuit implementation involving a single horizontal-cell-like feedback neuron - all despite their eyes having evolved independently since long before the emergence of image-forming vision in any animal. Here, the authors draw on Buchsbaum and Gottschalk’s ideas on efficient encoding (*1*) to suggest that like for zebrafish bipolar cells (*13*), the *Drosophila* R7/8 single and double zero-crossings can be conceptually matched with opsin-based primary and secondary colour axes, respectively. However, how this link would look like in practise for the encoding of spectral variance in natural light remained unclear. Here, we extend these theoretical links to directly show how like in zebrafish, *Drosophila* PC1 and PC2 are each well captured by a single receptor, while capturing PC3 requires possible opposing of multiple receptors downstream.

### Achromatic signaling

Natural scenes are generally dominated by achromatic over chromatic contrasts (*5*), and biased to mid- or long-wavelengths. Accordingly, an efficient achromatic encoder should approximate the resultant mid-/long-wavelength biased mean spectrum of light in a non-opponent manner - as is the case for both zebrafish red-cones (Fig. 2n) and for *Drosophila* yR8 photoreceptors (Fig. 6d). Here, the quality of the spectral match primarily impacts the maximal achievable signal-to-noise of the encoder, rather than its ability to encode brightness *per se* (*46, 47*). Accordingly, despite their minor respective mismatches compared to the mean of available light (see below), both zebrafish (Fig. 6a,b) and *Drosophila* implementations (Fig. 6d,e) capture PC1 well. For the same reason, also other non-opponent photoreceptors, such as *Drosophila* R1-6 (*48*) as well as vertebrate rods or “true” double-cones in many non-mammalian vertebrates, are generally thought to capture achromatic signals (*4*). However, in all these cases the presumed non-opponent nature at the level of their synaptic output *in vivo* remains to be confirmed.

In both zebrafish red-cones, and in *Drosophila* yR8, the largest mismatch to their natural environment’s PC1 was in the UV-range (Figs. 2n, 6d). Here, it is tempting to speculate that their low short-wavelength sensitivity is linked to a need to isolate behaviourally critical “general” achromatic signals from those that incur specifically in the UV-range. In the case of zebrafish, UV-specific signals carry key visuo-ecological relevance, in that they can report the presence of prey (*10*) – a rare feature that is unlikely to be captured in our scene-wide data of natural spectra (see also discussion on UV-signalling below).

Ultimately, the signals from red-cones must be read out by downstream circuits, in a manner that approximately preserves their spectral tuning. This could principally occur via a private-channel, as potentially provided by mixed-bipolar cells which in adults receive direct inputs only from red-cones and from rods (*49*). However, most zebrafish bipolar cells receive direct inputs from more than one cone type, presumably mixing their spectral signals. Nevertheless, a PC1-like signal does filter all the way to the brain where it forms the dominant Off-response (*47*).

### Primary chromatic signaling

In natural scenes, all spectral variance that is not captured by PC1 is chromatic, with any subsequent components capturing progressively smaller fractions of the remaining variance in a mutually orthogonal manner. Accordingly, PC2 and PC3 are maximally informative about primary and secondary spectral contrasts, respectively, while at the same time being uninformative both about brightness (i.e. PC1), or about each other. Here, we found that zebrafish green-cones (Fig. 2n), as well as *Drosophila* yR7 photoreceptors, both provide a good match to their respective environment’s PC2 (Fig. 6d). In the case of zebrafish, this match was close to perfect: When challenged with natural spectra, green-cones were highly informative about PC2, but uninformative about PC1 or PC3. Accordingly, like for red-cones (discussed above), the visual system would be well-served to read out the signal from green-cones in a private-line at least once so as to preserve this already efficient chromatic signal. Indeed, green-cones are anatomically the only cones in the zebrafish retina known to have such an arrangement: two of the more than twenty zebrafish bipolar cell “morpho-types”, both stratifying in the traditional “Off-stratum” of the inner plexiform layer (IPL), make exclusive contacts to green cones (*49*). Potentially in agreement, we previously identified a small but well-defined population of singly colour-opponent bipolar cell responses in this part of the IPL (*13*).

### Further chromatic signaling

Beyond PCs 1 and 2, most of the remaining spectral variance was captured by PC3, which presents a triphasic spectral response with two zero crossings. However, neither of the remaining blue- and UV-cones exhibited such a tuning. Of these, blue-but not UV-cones were strongly opponent, nevertheless suggesting their important role in spectral processing. Accordingly, we explored why blue-cones did not directly capture PC3. For this, we returned to our horizontal cell model, this time immediately optimising red-green- and blue-cones to match PC1, PC2 and PC3, respectively. To complete the model, UV-optimisation was left unchanged to again target its own *in vivo* tuning function. Using this strategy, it was possible to produce only weakly distorted red-, green- and UV-cone spectra. However, the model failed to directly capture PC3 using blue-cones, and the mild relative distortion of green-cone spectral tuning was sufficient to noticeably degrade their ability to capture PC2 (Fig. S6h-k). This tentatively suggests that the specific connectivity of the outer retina, constrained by the four principal zebrafish cone-opsins, is poorly suited to additionally produce a PC3-like spectral response.

Nevertheless, blue-cones did exhibit a single zero crossing that differed from that of green-cones, meaning that two zero crossing could be readily achieved in a linear model that opposed green- and blue-cone signals (Fig. S5). We showed that such an arrangement would at least in theory allow building a spectral filter which closely captures PC3 while producing only poorly correlated responses to PC1 and PC2. Intriguingly, such a PC3-like filter is in fact observed at the level of the brain, which mainly opposes UV- and Red- “On” signals with spectrally intermediate blue/green “Off” signals (*47*). However, how this brain response is set-up at the level of the retina, remains unclear. Finally, a PC3-like signal could also be achieved in *Drosophila* by opposing their two mid-wavelength sensitive yR7 and pR8 photoreceptors, however in this case the best match was achieved when in addition recruiting the more broadly tuned yR8 alongside yR7 (Fig. S6d,f).

### A private channel for detecting UV-signals?

Remarkably, unlike red-green- or blue-cones, the final output of zebrafish UV-cones appeared to not be central to support dominant achromatic nor chromatic processing. In fact, UV-cones also use a nearly UV-exclusive horizontal cell (H3, Figs. 3,4) (*30, 33*), likely for temporal tuning (*10*), while barely contributing to the signals of H1 and H2 (Fig. 4d). Accordingly, outer retinal UV-circuits appear to approximately signal in isolation from those of the remaining cones. Similarly, direct contributions from the UV-sensitive pR7 photoreceptors were also not required to approximate the first three PCs that emerge from the natural spectral world of *Drosophila* (Fig. 6d,e). In both cases, these photoreceptors contrasted their strong, short-wavelength exclusive response with weaker opposition at most other wavelengths. From here, it is tempting to speculate that these UV-systems may serve to *detect*, rather than necessarily to *spectrally contrast*, the presence of strongly UV-biased objects against a “naturally-grey” background. Such a detector would be invaluable for reporting the presence of the UV-bright single-celled microorganisms when illuminated by the sun, which larval zebrafish feed on (*10*). To our knowledge, a similarly specific visuo-ecological purpose of UV-vision in *Drosophila* remains unknown. More generally, UV-light can be highly informative about edges in space, as it tends to accentuate objects’ silhouettes against bright backgrounds (*50*– *52*).

In zebrafish, previous work has highlighted a key role of UV-vision across the retina and brain leading to behaviour (*10, 13, 24, 47, 53, 54*). Most notably, the retina’s acute zone (*25*) is dominated by UV-sensitive circuits (*13*). Here, most bipolar cell terminals respond primarily to UV-stimulation, and only some in addition respond to other wavelengths (*13*) – a general pattern that is recapitulated also at the level of the retinal ganglion cells (*24*) to drive a strong UV-response in the brain (*47, 55, 56*) which filters all the way to spinal circuits (*56, 57*). Nevertheless, despite this profound functional dominance, no anatomical study has reported the presence of UV-cone-dedicated bipolar cells, as for example in the case of green-cones (*49*) (see above). While it remains unknown if such connectivity specifically exists in the acute zone, it seems clear that more broadly across the retina, the signals from UV-cones are mixed with those of other cones. How this connectivity serves to support the diverse visuo-ecological needs of zebrafish UV-vision will be important to address in the future.

### Regional differences in cone spectral tuning

Unlike many other aspects of larval zebrafish retinal structure (*13, 24*–*26, 58*) and function (*10, 13, 24*), the spectral tuning of zebrafish cones was remarkably eye-region invariant (Fig. S1g). Nevertheless, small but significant regional variations were observed in all cone-types. Of these, the most striking differences occurred in red- and green-, and to a smaller extent also in UV-cones. Red-cones, and to a weaker extend also other cones, exhibited relatively narrowed tuning ventrally, and broadened tunings dorsally. These differences might help keeping cones within operational range despite the large difference in absolute amount light driving them: bright direct skylight versus dimmer reflected light from below, respectively. Next, amongst green-cones, the acute-zone exhibited the strongest short-wavelength response, resulting in a long-wavelength shift in their zero crossing. This finding is conceptually in line with an increase in absolute light sensitivity amongst UV-cones in this part of the eye (*10*), however a possible visuo-ecological purpose of this shift remains to be established. Finally, mid-wavelength opponency amongst UV-cones was strongest in the AZ and ventrally, which may be linked to the behavioural need to contrast UV-bright prey against a spectrally intermediate but bright background in the upper-frontal parts of visual space (*10, 59*). In contrast, larval zebrafish rarely pursue prey below or behind them (*59, 60*), as surveyed by dorsal and nasal UV-cones, respectively.

Finally, we wondered how eye-region differences in cone-tunings might be achieved at the level of outer retinal circuits. To explore this, we again returned to our horizontal cell model, this time fitting it individually to only the subsets of recordings from each of the four regions. This revealed that the same anatomically established maximal connectivity matrix (Figs. 3,4) served well to produce any of these regional differences by minimally shifting their relative weights (Table S1). Accordingly, it seems likely that the same principal horizontal cell network produces these regional variations in tuning based on minor rebalancing of its relative input strengths.

## METHODS

### RESOURCE AVAILABILITY

#### Lead Contact

Further information and requests for resources and reagents should be directed to and will be fulfilled by the Lead Contact, Tom Baden.

#### Material Availability

Plasmids pBH-opn1sw2-SyGCaMP6f-pA, pBH-LCRhsp70l-SyGCaMP6f-pA, pBH-thrb-SyGCaMP6f-pA, pBH-cx55.5-nlsTrpR-pA, pBH-tUAS-ASAP3-pA and transgenic lines *Tg(opn1sw2:SyGCaMP6f), Tg(LCRhsp70l:SyGCaMP6f), Tg(thrb:SyGCaMP6f), Tg(cx55*.*5:nlsTrpR,tUAS:ASAP3)*, generated in this study, are available upon request to the lead contact.

#### Data and Code Availability

Pre-processed functional 2-photon imaging data, natural imaging data, EM-data, HC circuit modelling data, and associated summary statistics will be made freely available via the relevant links on http://www.badenlab.org/resources and http://www.retinal-functomics.net. Code for the model and the statistical analysis of the experimental data is available on Github (https://github.com/berenslab/cone_colour_tuning). Natural imaging datasets were published previously as part of (*12, 13*).

## EXPERIMENTAL MODEL AND SUBJECT DETAILS

### Animals

All procedures were performed in accordance with the UK Animals (Scientific Procedures) act 1986 and approved by the animal welfare committee of the University of Sussex. For all experiments, we used *7-8 days post fertilization* (*dpf*) zebrafish (Danio rerio) larvae. The following previously published transgenic lines were used: *Tg(opn1sw1:nfsBmCherry)*(*33*), *Tg(opn1sw1:GFP)*(*61*), *Tg(opn1sw2:mCherry)*(*62*), *Tg(thrb:Tomato)*(*63*). In addition, *Tg(opn1sw2:SyGCaMP6f)*, and *Tg(LCRhsp70l:SyGCaMP6f), Tg(thrb:SyGCaMP6f)*, lines were generated by injecting pBH-opn1sw2-SyGCaMP6f-pA, pBH-LCRhsp70l-SyGCaMP6f-pA, or pBH-thrb-SyGCaMP6f-pA. *Tg(cx55*.*5:nlsTrpR,tUAS:ASAP3)*, line was generated by co-injecting pBH-cx55.5-nlsTrpR-pA and pBH-tUAS-ASAP3-pA plasmids into single-cell stage eggs. Injected fish were out-crossed with wild-type fish to screen for founders. Positive progenies were raised to establish transgenic lines.

All plasmids were made using the Gateway system (ThermoFisher, 12538120) with combinations of entry and destination plasmids as follows: pBH-opn1sw2-SyGCaMP6f-pA: pBH (*33*) and p5E-opn1sw2 (*33*), pME-SyGCaMP6f (Yoshimatsu et al., 2020), p3E-pA(*64*); pBH-LCRhsp70l-SyGCaMP6f-pA: pBH and p5E-LCRhsp70l, pME-SyGCaMP6f, p3E-pA; pBH-thrb-SyGCaMP6f-pA: pBH and p5E-1.8thrb (*63*), pME-SyGCaMP6f, p3E-3.2thrb (*63*); pBH-tUAS-ASAP3-pA: pBH and p5E-tUAS (*65*), pME-ASAP3, p3E-pA. Plasmid p5E-LCRhsp70l was generated by inserting a polymerase chain reaction (PCR)-amplified Locus Control Region (LCR) for green opsins (RH2-1 to 2-4) (*66*) into pME plasmid and subsequently inserting a PCR amplified zebrafish 0.6 kb hsp70l gene promoter region (*67*) downstream of LCR. pME-ASAP3 was made by inserting a PCR amplified ASAP2s fragment (*68*) and subsequently introducing L146G, S147T, N149R, S150G and H151D mutations (*37*) in pME plasmid.

Animals were housed under a standard 14:10 day/night rhythm and fed three times a day. Animals were grown in 0.1 mM 1-phenyl-2-thiourea (Sigma, P7629) from 1 *dpf* to prevent melanogenesis. For 2-photon *in-vivo* imaging, zebrafish larvae were immobilised in 2% low melting point agarose (Fisher Scientific, BP1360-100), placed on a glass coverslip and submerged in fish water. Eye movements were prevented by injection of a-bungarotoxin (1 nL of 2 mg/ml; Tocris, Cat: 2133) into the ocular muscles behind the eye. For some experiments, CNQX (∼0.5 pl, 2 mM, Tocris, Cat: 1045) or meclofenamic acid sodium salt (MFA) (∼0.5 pl, 5 mM, Sigma, Cat: M4531) in artificial cerebro-spinal fluid (aCSF) was injected into the eye.

## METHOD DETAILS

### Light Stimulation

With fish mounted on their side with one eye facing upwards towards the objective, light stimulation was delivered as full-field flashes from a spectrally broad liquid waveguide with a low NA (0.59, 77555 Newport), positioned next to the objective at ∼45°. To image different regions in the eye, the fish was rotated each time to best illuminate the relevant patch of photoreceptors given this stimulator-geometry. The other end of the waveguide was positioned behind a collimator-focussing lens complex (Thorlabs, ACL25416U-A, LD4103) which collected the light from a diffraction grating that was illuminated by 14 spectrally distinct light-emitting diodes (LEDs) (details on LEDs below). Following the an earlier design (*14*), the specific wavelength and relative angle of each LED to the diffraction grating defined the spectrum of light collected by the collimator to be ultimately delivered to the fish’s eye, according to:

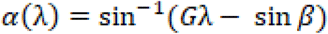

where α is the angle of light incident to the diffraction grating, λ the wavelength (in nm), β the first order diffraction exit angle and G the diffraction grating’s groove density. Moreover, each LED was individually collimated (Signal Construct SML 1089 - LT-0454) and attached to a rail (Thorlabs, XE25L450/M; XE25L225/M) by a 3D printed holder (available at https://github.com/BadenLab/HyperspectralStimulator).

An Arduino Due (Arduino) and LED driver (Adafruit TCL5947) were used to control and drive the LEDs, respectively. Each LED could be individually controlled, with brightness defined via 12-bit depth pulse-width-modulation (PWM). To time-separate scanning and stimulating epochs, a global “blanking” signal was used to switch off all LEDs during 2P scanning but enable them during the retrace, at line-rate of 500 Hz (see also (*15, 16*). The stimulator code is available at https://github.com/BadenLab/HyperspectralStimulator).

LEDs used were: Multicomp Pro: MCL053RHC, Newark: C503B-RAN-CZ0C0AA1, Roithner: B5-435-30S, Broadcom: HLMP-EL1G-130DD, Roithner: LED-545-01, TT Electronics: OVLGC0C6B9, Roithner: LED-490-06, Newark: SSL-LX5093USBC, Roithner: LED450-03, VL430-5-1, LED405-03V, VL380-5-15, XSL-360-5E. Effective LED peak spectra as measured at the sample plane were, respectively (in nm): 655, 635, 622, 592, 550, 516, 501, 464, 448, 427, 407, 381, 361, 360 nm. Their maximal power outputs were, respectively (in µW): 1.31, 1.06, 0.96, 0.62, 1.26, 3.43, 1.47, 0.44, 3.67, 0.91, 0.24, 0.23, 0.04, 0.20. From here, the first ten LEDs (655 – 427 nm) were adjusted to 0.44 µW, while the four UV-range LEDs were set to a reduced power of 0.2 µW (407, 381, 360 nm) or 0.04 µW (361 nm). This relative power reduction in the UV-range was used as a compromise between presenting similar power stimulation across all LEDs, while at the same time ameliorating response-saturation in the UV-range as a result of the UV-cones’ disproportionately high light sensitivity (*10, 24*). In this regard, we took advantage of the strong spectral overlap between the two shortest-wavelength LEDs (360, 361 nm) to probe this wavelength range at two intensities (0.2 and 0.04 µW, respectively).

From here, all spectral tuning functions were based on the responses to the 13 spectrally distinct LEDs, excluding the response to low-power 361 nm LED. This strategy yielded biologically highly plausible spectral sensitivity functions in all cones that closely resembled their underlying opsin’s tuning when pharmacologically isolated from horizontal cells (Fig. 2b). Nevertheless, UV-cones weakly but consistently under-shot their opsin template at the shortest tested wavelength (360 nm), hinting that they may have approached their saturation point at this wavelength and power. In agreement, the 0.04 µW 361 nm LED elicited only mildly lower response-amplitudes in UV-cones compared to the 0.2 µW 360 nm LED (R_low_ = 0.88±0.14; R_high_ = 0.96±0.06, errors in SD; difference p <<0.001 Wilcoxon signed-rank test). In contrast, all other cones responded much more weakly to the low power UV-LED: Blue-cone (R_low_ = 0.35±0.16; R_high_ = 0.67±0.21); green-cone (R_low_ = −0.12±0.24; R_high_ = 0.09±0.32); red-cone (R_low_ = −0.02±0.27; R_high_ = 0.21±0.27; all low-high pairs p << 0.001) suggesting that these cones were not near their UV-saturation points.

Together, it therefore remains possible that measured cone-tuning functions relatively underestimate UV-components, however this effect is likely to be very small in non-UV-cones that dominate “traditional” colour vision in zebrafish (Discussion). The exact slope of the cones’ UV-response also had negligible impact on their relative matches with PCs or their contributions to the HC-network (not shown), in line with an only weak interdependence of the outer retina’s UV-versus red-/green-/blue-cone systems (see Discussion).

### 2-photon calcium and voltage imaging

All 2-photon imaging was performed on a MOM-type 2-photon microscope (designed by W. Denk, MPI, Martinsried; purchased through Sutter Instruments/Science Products) equipped with a mode-locked Ti:Sapphire laser (Chameleon Vision-S, Coherent) tuned to 960 nm for SyGCaMP6f and ASAP3 imaging. To measure HC tuning functions, we first expressed GCaMP6f in HCs. However, while we observed strong light-driven calcium responses at their dendritic tips, adjacent to cone terminals and thus indicative of local processing, we did not observe robust calcium responses in the HC soma (as a proxy of global processing). This lack of somatic calcium responses could be due to a putative lack of voltage-gated calcium channels in larval HC somata (unlike e.g. in adult mouse (*69*)). Instead, we therefore measured voltage responses using the genetically encoded voltage sensor, ASAP3(*37*), which presumably also gave a more direct readout of HC global function. We used two fluorescence detection channels for SyGCaMP6f/ASAP3 (F48×573, AHF/Chroma) and mCherry (F39×628, AHF/Chroma), and a water immersion objective (W Plan-Apochromat 20x/1,0 DIC M27, Zeiss). For image acquisition, we used custom-written software (ScanM, by M. Mueller, MPI, Martinsried and T. Euler, CIN, Tuebingen) running under IGOR pro 6.3 for Windows (Wavemetrics). Recording configurations were as follows: UV-cone SyGCaMP6f 128×128 pixels (2 ms per line, 3.9 Hz) or 256×256 pixels (2 ms per line, 1.95 Hz); all other cones SyGCaMP6f and horizontal cell ASAP3 256×256 pixels (2 ms per line, 1.95 Hz).

### Pre-processing and extraction of response amplitudes of 2-photon data

Regions of interest (ROIs), corresponding to individual presynaptic cone terminals were defined automatically based on local thresholding of the recording stack’s standard deviation (s.d., typically > 25) projection over time, followed by filtering for size and shape using custom written scripts running under IGOR Pro 6.3 (Wavemetrics), as used previously in (Yoshimatsu et al., 2020). Specifically, only ellipsoidal ROIs (<150% elongation) of size 2-5 μm^2^ were further analyzed. For ASAP3 recordings, ROIs were manually placed to follow the shape of individual HC somata. Calcium or voltage traces for each ROI were extracted and z-normalized based on the time interval 1-6 s at the beginning of recordings prior to presentation of systematic light stimulation. A stimulus time marker embedded in the recording data served to align the traces relative to the visual stimulus with a temporal precision of 2 ms.

Following the approach used in (*70*), a quality criterium (QC) of how well a cell responded to a stimulus were computed as

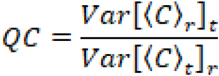

where C is the T by R response matrix (time samples by stimulus repetitions) and ⟨ ⟩x and *Var*[]x denote the mean and variance across the indicated dimension, respectively. If all trials are identical such that the mean response is a perfect representative of the response, QC is equal to 1. If all trials are random with fixed variance, QC is equal to 1/R. For further analysis, we used only cells that responded well to the stimulus (QC >0.4 for SyGCaMP6f or >0.32 for ASAP3) (see also SFig. 2b)

After filtering out poorly responsive cells using QC, outliners were removed using principal component analysis. Because in all cone types, PC1 explained >80% variance of the data, we computed the loading values of the principal component 1 of cone tuning function within each cone type and defined outliners as the cones with PC1 loading below 1.25 times the length of the 97 percentile departure from the mean.

To extract response amplitudes to each stimulus wavelength, an exponential curve was fit to the entire rising (or falling, for hyperpolarising responses) phase during each stimulus presentation, with the maximum value of the fitted curve was taken as the response amplitude. Because cones are intrinsically “Off-cells” (i.e. hyperpolarize to light) we then sign-inverted extracted amplitude values such that Off-responses would yield positive amplitude readings, and vice versa for On-responses. However, for voltage imaging, because ASAP3 fluorescence intensity increases as cells hyperpolarize, we preserved the polarity of the response amplitudes.

### Immunostaining and confocal imaging

Larval zebrafish (*7-8 dpf*) were euthanised by tricane overdose and then fixed in 4% paraformaldehyde (PFA, Agar Scientific, AGR1026) in PBS for 30 min at room temperature. After three washes in PBS, whole eyes were enucleated and the cornea was removed by hand using the tip of a 30 G needle. Dissected and fixed samples were treated with PBS containing 0.5% TritonX-100 (Sigma, X100) for at least 10 mins and up to 1 day, followed by the addition of primary antibodies. After 3-5 days incubation at 4°C, samples were washed three times with PBS 0.5% TritonX-100 solution and treated with secondary antibodies. After one day incubation, samples were mounted in 1% agar in PBS on a cover slip and subsequently PBS was replaced with mounting media (VectaShield, H-1000) for imaging. For HC imaging (Fig. S6c-f), the retina was flat-mounted with the photoreceptors facing to the cover slip. For cone side-view imaging (Fig. S6a), the lens was kept attached the retina to maintain the spherical shape of the retina, with the whole “retina-ball” mounted with the lens side facing to the cover slip. All presented data was imaged in the acute zone.

Primary antibodies were zpr-1 antibody (mouse, 1:100, ZIRC). Secondary antibodies were DyLight647 anti-mouse (Donkey, 1:500, Jackson Immunoresearch Laboratories). Confocal image stacks were taken on a TSC SP8 (Leica) with a 63x oil immersion objective (HC PL APO CS2, Leica). Typical voxel size was 90 nm and 0.5 μm in xy and z, respectively. Contrast, brightness and pseudo-colour were adjusted for display in Fiji (NIH).

To sparsely label HCs, plasmids pCx55.5:Gal4 and pUAS:MYFP were co-injected into one-cell stage eggs (*71*).

### UV-cone ablation

Larval zebrafish were immersed in fish water containing 10 mM Metronidazole (Met) for 2 hours to ablate nfsB-expressing UV-cones. Following Met treatment, zebrafish were transferred into fish water without Met and fed regularly until used for two-photon imaging.

### Electron-microscopy data acquisition, reconstruction and annotation

A larval zebrafish (*8 dpf*) was euthanised by tricane overdose and then a small incision on a cornea was made using 30G needle in a fixative solution containing 4% glutaraldehyde (AGR1312, Agar Scientific,) in 0.12M cacodylate buffer, pH 7.4. The tissue was immediately transferred into a 1.5 ml tube with the fixative, centrifuged at 3,000 rpm for 3 min, and further fixed in the fixative over-night on a shaker at room temperature. Subsequently, the tissue was washed 3 times in 0.12M cacodylate buffer, pH7.4 and incubated in a solution containing 1.5% potassium ferrocyanide and 2% osmium tetroxide (OsO4) in 0.1M cacodylate buffer (0.66% lead in 0.03M aspartic acid, pH 5.5) for 1 hour. After washing, the tissue was placed in a freshly made thiocarbohydrazide solution (0.1g TCH in 10 ml double-distilled H20 heated to 600 C for 1 h) for 20 min at room temperature (RT). After another rinse, at RT, the tissue was incubated in 2% OsO4 for 30 min at RT. The samples were rinsed again and stained *en bloc* in 1% uranyl acetate overnight at 40 C, washed and stained with Walton’s lead aspartate for 30 min. After a final wash, the retinal pieces were dehydrated in a graded ice-cold alcohol series, and placed in propylene oxide at RT for 10 min. Finally, the sample was embedded in Durcupan resin. Semi-thin sections (0.5 −1 µm thick) were cut and stained with toluidine blue, until the fiducial marks (box) in the GCL appeared. The block was then trimmed and mounted in a Serial-blockface scanning electron microscope (GATAN/Zeiss, 3View). Serial sections were cut at 50 nm thickness and imaged at an xy resolution of 5 nm. Two tiles, each about 40 µm x 40 µm with an overlap of about 10%, covering the entire photoreceptor and horizontal cell layers in a side view at the acute zone were obtained. The image stacks were concatenated and aligned using TrackEM (NIH). The HCs and cones were traced or painted using the tracing and painting tools in TrackEM2 (*72*).

### Clustering of HCs in EM and Confocal data

To validate the ad hoc group assignment based on UV contacts (HC area) and R/G contacts for the electron microscopy (Fig. 3h,i) and confocal data (Fig. S3g) we used Mixture of Gaussian (MoG) clustering on all extracted features. These features (area size, number of contacts to R/G, B, U, for EM and area size, tip density, number of contacts to R, G, B/U for CM) were z-normalized and clustered in the same framework as the HC recordings (see below). The MoG clusters did coincide with the ad hoc group assignment.

### Opsin Templates and log transforms

For the log-transformed opsin templates (Fig. 1f, 2b) we assumed a baseline activation (represented by *b* in Eq. 2) and fit a linear transformation to take the arbitrary scaling of the recordings into account. We then optimized the function *f*_*a,b,c*_ to minimize the mean squared error (MSE) between *f*_*a,b,c*_ (*opsin*) and the data of the HC block condition for each cone type:

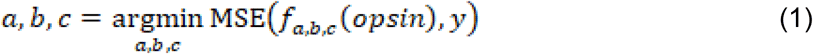

where y is the mean of the HC block condition and f is the function

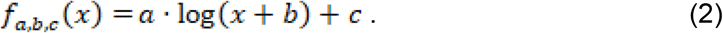

For the optimization we used the python package scipy.optimize.minimize (version 1.4.1). The inverse of this procedure is shown in Fig. S4a, where the mean of HC block condition is fitted in the same way to the opsin curves of each cone with the function:

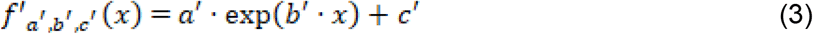

The data distribution (25 and 50 and 75 percentiles) is then calculated by passing each individual HC block recording through the optimized function *f*′.

### Model of cone and HC interaction

We modelled cone-HC interactions as a linear model and included the established (Fig. 3k) connectivity pattern for the three types of HC as a (3×4) connectivity matrix *W* where *w*_*ij*_ indicates connection strength from cone type *j* to HC type *i*. Further, we assumed the feedback strength per connection of each HC type to be constant for all cones and defined it as a diagonal matrix *A*. To compute the effective feedback, this matrix is then weighted by the relative connection strength per cone and HC, represented in a (4×3) matrix *F* with 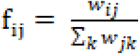. This represents the strength from HC type *j* to cone of type *i*. Hereby we assume a symmetric connectivity pattern which is justified by the symmetrical cone mosaic in zebrafish. With these definitions, we can formulate the model recurrently as following:

The inputs to the HCs is defined as

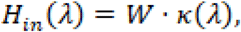

where *k*(*λ*)represents the raw activity in the synapse, which still has to be shifted according to the baseline. The summed outputs of the HCs are computed as

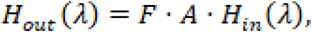

Finally, the raw activity in the synapses is computed as

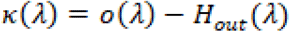

where *o*(*λ*) represents the wavelength dependent opsin activation.

The same formulas hold for computing the baseline of the cones, for which *o*(*λ*) was set to **1**, which accords to the applied normalization on the recorded data. The final output of the model are the tuning curves *k* shifted to the cone specific baselines and normalized.

The same normalization procedure was applied to the shown HC spectra, which are the normalized spectra *H*_*in*_(*λ*).

In the reduced models, in which we only included specific types of HCs, we set the corresponding entries in the weight matrix *W* to zero but did not change the model otherwise.

#### Model input

To extract the cone tuning curves from the experimental data for the model, we computed the mean amplitude of each bright and dark three seconds interval but excluded in each interval the first second as adaption time. We then took for every individual trace the difference of each bright interval to its preceding dark interval based on these means. Finally, we averaged over these values for each cone type and experimental condition and, by assuming smooth tuning functions, interpolated (using the scipy function *scipy*.*interpolate*.*interp1d*) the data to an equidistant resolution of 1nm.

As input to our model we took the normalized traces of the blocked HC condition. This normalization can be interpreted as a maximal dark current of 1 and a minimal current of 0 during activation. The input acted as “opsin-sensitivity” curves *o*(*λ*) of the cones. We decided to use these curves instead of the theoretical available opsin tuning curves since we have a pure linear model and as shown in Fig. 2b these traces are a good proxy for the log-transformed opsin templates, which is the effective activation for this linear model. All spectral tuning curves of the cones were normalized to have a maximal absolute value of one.

#### Fitting procedure

We used the Sequential Neural Posterior Estimation method (also called SNPE-B) described in (*36*) (code available at https://github.com/mackelab/delfi, version: 0.5.1) with small modifications which were already applied in (*73*) to fit our model.

In brief, SNPE-B draws parameters {*θ*_*i*_}_*i*∈*I*_ over several rounds r=1,…R from a (proposal) prior 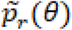 and evaluates the model for these parameters. For the evaluations *e*_*i*_*(θ*_*i*_) the discrepancy function *x*_*i*_(*e*_*i*_) = *D*(*e*_*i*_) is computed and a mixture density network (MDN) q_Φ_(*θx*) is trained on the data pairs {(θ_i_,x_i_)}_*i*∈*I*_. The posterior *p*_*r*_*(θ*|*x*_0_) is then calculated as *q*_*Φ*_(*θ*|*x = x*_0_) and used as a new proposal prior in the next sampling round: 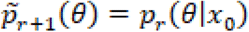. We took the MSE between model output and the data as discrepancy function. This implies *x*_0_ = 0, but as our data is noisy, our model cannot get to a MSE of zero. This would mean, that the MDN has to extrapolate to unreached discrepancy values, which could lead to an unstable behaviour. As a consequence, we took as *x*_0_ the 0.01-percentile of {*x*_*i*_}_*i*∈*I*_ in each round. This evaluation of *q*_*Φ*_(*θ*|*x = x*_0_) can be understood as the posterior over the parameters for the “best possible” model evaluations. Testing for different percentiles in a reasonable range did not change the results. We took the same approach for setting an adaptive bandwidth for the kernel (see also (*73*)). As for a few models the posteriors became slightly worse after some rounds, we compared post-hoc the posterior distributions of each round and took the one with the smallest 1-percentile of its samples.

We ran SNPE-B over five rounds, with 200,000 samples per round. The prior was a multivariate normal distribution with mean 1_*n*_ and covariance 0.25· *Id*_*n*_, where *n* is the number of model parameters, ranging from 11 (all HCs) to 3 (only H2). We chose three Gaussian components for the MoG and a MDN with two hidden layers with 100 nodes each. In each round the network was trained for 600 epochs with a minimum batch size of 500 and continuous learning started in round two. To let the MDN focus on regions of low discrepancy, we used a combined Uniform-Half-Gaussian kernel which was constant 1 up to *x*_0_ and decayed then as a half Gaussian. The scale of the Half-Gaussian part was in each round chosen as the 20-percentile of the discrepancy values. For the presented tuning curves 100,000 samples were drawn from the final posterior and the model evaluated.

### HC clustering based on spectral tuning

To identify functional clusters we used a Mixture of Gaussians model (sklearn.mixture.GaussianMixture, version 0.21.2) with three components and diagonal covariance matrices on the pre-processed tuning curves (n = 86) which were additionally normalized to have maximal value of one. Aiming for a stable clustering, we ran the algorithm 1,000 times with different random seeds and chose the ones with the smallest BIC and under these chose the partition which appeared most often. The different runs did not change the general shape of the cluster means, but the specific assignment was variable for some traces. With this procedure we got a partition with n = 12, 19, 55 elements, which were allocated to the known functional tunings for HCs of adult zebrafish (*30, 34*).

### Natural Imaging Data Analysis

The hyperspectral data were element-wise multiplied with a deuterium light source derived correction curve [S.x]. The data were restricted to the domain of 360-650 nm and z-normalised within a given scan. Here, the long-wavelength end of the domain was decided based on the long-wavelength opsin absorption curve; the short-wavelength end was dictated by the sensitivity of the spectrometer. The hyperspectral PCs were obtained using the scitkit-learn 0.22.1 implementation of the Principal Component Analysis algorithm. Only the first three components are displayed.

Hyperspectral measurement points were spatially aligned within the scan according to the scan raster (see (*12, 13*) for details). Pixel brightness is the projection of a given PC, or mean of the convolution with the opsin absorption or the observed cone response curves respectively. Presented images were smoothed using a Gaussian filter (sigma = 2px). Sum of Squares difference was taken between pairs of z-normalised images as well as their negatives. The lowest Sum of Squares (=highest correlation, either with the original or the negative) is displayed. Smoothing did not significantly affect this measure.

To statistically compare scene reconstructions by different sets of tuning functions (Fig. S6a-c), we used two parallel strategies. First, we computed the correlation coefficient between reconstructions by the different channels (e.g. *in vivo* red cone vs. green cone) as indicated for each of n = 30 scenes, thus yielding 30 correlation coefficients for each combination of channels in each condition. Amongst each comparison we then computed the mean and SD, as shown.

Second, to capture the multivariate dependence directly, we computed the mutual information under Gaussian assumption, *MI* = Σ_*i*_ *h*(*x*_*i*_)−*h*(*x*)∼log det[2 *π e C*], where *C* is the correlation matrix of the scene representations in the different channels (e.g. 4×4 *in vivo*: red-, green-, blue-, UV-cone). As the diagonal of *C* is constant and equal to, the mutual information is proportional to the latter quantity. We normalized this quantity by the mutual information of the opsin set of tuning functions.

### Linking opsin- and photoreceptor-spectra to principal components

Measured *in vivo* spectra of cones and their underlying log-transformed opsin templates (Fig. 1f) were linearly combined to provide least-squares fits to the respective underwater spectral PCs (Figs. 5n, 6d, Figs. S5,6). The same procedure was also used to match *Drosophila* R7/8 spectra (Fig. 6c, from (*11*)) to the PCs that emerged from natural distribution of light above the water. Next, to compare the expected responses of *in vivo* photoreceptors, their linear combinations (in case of PC3, see below), as well as their respective log-opsin constructs to natural light, individual natural light pixel spectra (n = 30,000) were multiplied with the respective sensitivity curves. In each case, pixel-spectra were first z-normalised within the scene, and products were summed over all wavelengths. This procedure produced ‘responses’ (Fig. 6a), which were plotted against the respective loadings of each spectrum onto PC1, PC2 and PC3 (in rows 1, 2 and 3, respectively). From here, scene-wise summary statistics were computed based on Spearman correlation coefficients (Fig. 6b,e).

To arrive at *in vivo* photoreceptor combinations that best approximated PC3s zebrafish: Fig. S5a-c, Drosophila: Fig. S6e,f), we assessed the spectral matches to them by several plausible linear combinations of *in-vivo* photoreceptor tunings based on least squares. In both cases, the best fits required opposing the two spectrally intermediate receptors. For zebrafish, this “GB-fit” performed as well as any combination of more complex fits that in addition used red- or UV-cones, so we used this simplest GB-fit for further analysis. In case of *Drosophila*, best performance required also adding the long-wavelength sensitive receptor to yield an yR8+yR7-pR8 axis (short: “yyp8”). In each case, performance as shown in Figs. S5c and Fig. S6f (top) was evaluated based on the mean scene-wise Spearman correlation coefficient between the resultant spectral axis, as described above. The weights needed to build these PC3-like tunings based on photoreceptor types are plotted below as abs(max)-normalised for better comparison.

## QUANTIFICATION AND STATISTICAL ANALYSIS

### Statistics

No statistical methods were used to predetermine sample size. Owing to the exploratory nature of our study, we did not use randomization or blinding.

We used Generalized Additive Models (GAMs) to analyse the relationships between wavelength and cone activity under different experimental conditions (Fig. 2d-h, Fig. S2). GAMs can be understood as an extension to the generalized linear model by allowing linear predictors, which depend on smooth functions of the underlying variables (*74*). We used the mgcv-package (version 1.8-31) in R on an Ubuntu 16.04.6 LTS workstation with default parameters. We modelled the dependence of the variable of interest as a smooth term with 13 degrees of freedom. The models explained ∼59-82% of the deviance. Statistical significance for differences between the dependence of activation in the different experimental conditions were obtained using the plot_diff function of the itsadug-package for R (version 2.3). Significance of opponency (Fig. S1g) and zero crossings of the tuning curves (Fig. 1f, Fig. S1g) were also calculated based on GAMs with “zone” as an additional predictive variable and grouping where applicable.

## Acknowledgements

We thank Sharm Knecht and Rachel Wong for EM volume acquisition. We thank Thomas Euler for critical feedback. The authors would also like to acknowledge support from the FENS-Kavli Network of Excellence and the EMBO YIP.

## Author contributions

TY, PBa and TB designed the study, with input from CS, FKJ and PBe. TY generated novel lines and performed 2-photon data collection and pre-processing. TY also performed anatomical imaging and EM-tracing. TY and CS analysed anatomical data. PBa built the light-stimulator with input from FKJ. PBa and TB performed natural imaging data analysis, with input from PBe. CS performed computational modelling of the HC-cone circuit with input from PBe. CS analysed voltage recordings with input from PBe. CS, TY, PBa and TB performed general statistical analyses, with help from PBe. FSP provided early access to ASAP plasmids. TB wrote the manuscript with inputs from all authors.

