## Supplemental Figures and Table for "Ancestral circuits for vertebrate colour vision emerge at the first retinal synapse"

Yoshimatsu et al.

### SUPPLEMENTAL MATERIALS

**6 Supplemental Figures**

**1 Supplemental Table**

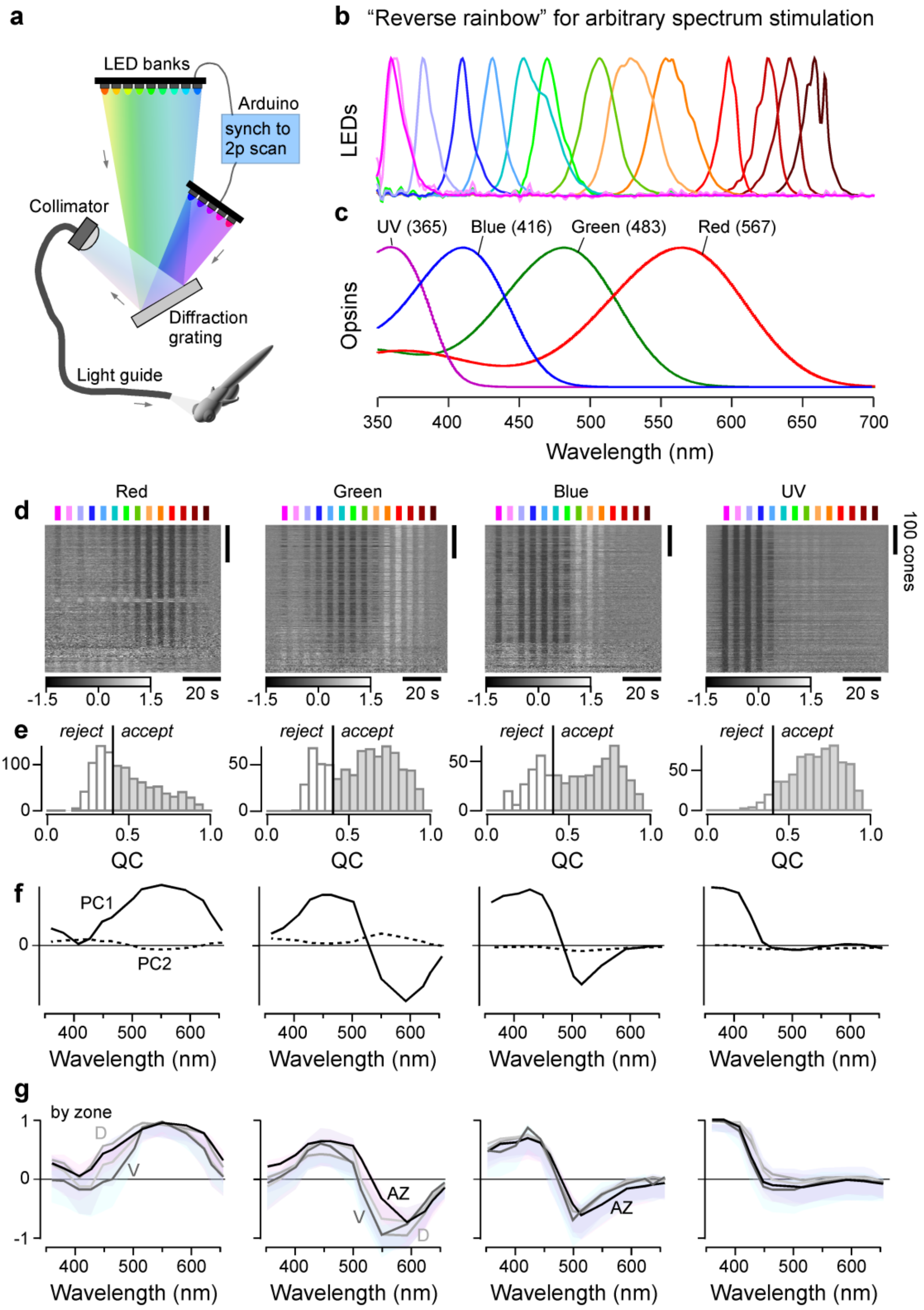


***Figure S1 | Hyperspectral stimulation under 2-photon and quality filtering of cone-recordings (related to Figure 1). a**, Schematic of the stimulator, inspired by (14): Two banks holding a total of 14 spectrally distinct and collimated LEDs are pointed at a diffraction grating. A spectrally narrowed fraction of each LED's light is then further collimated into a light guide and presented as full-field to the zebrafish under the 2P microscope. To prevent spectral cross-talk, the LEDs are time-interleaved with the 2P scan (15). **b**, Peak normalised spectra of the 14 LEDs measured at the sample plane. LED Powers were presented to be equal across the full spectrum, with exception of the four short-wavelength ones which were relatively attenuated to ~50% power to ameliorate UV-saturation (Methods). Moreover, the strong spectral overlap in the two shortest wavelength LEDs was exploited as an internal UV-saturation control, by tuning the second LED to ~10% output power (Methods). **c**, Govadovskii-templates of opsin absorption spectra for the four zebrafish cones(77, 78). **d,e** heatmaps of all cone recordings of each type as indicated (n = 1,659), ranked by quality criterion (b) (QC, Methods) from these*

*recordings, with a cut-off at QC = 0.4, as indicated. **f**, First principal components from tunings extracted from all ROIs of a given cone type (PC1 and 2 are shown). In all cone types, PC1 explains >80% of the variance in the data. Thus, we used PC1 loadings to remove outliers (Methods). **g**, Means±SD for all ROIs that passed QC and outliner filtering, segregated by recording region (acute zone (AZ), black; dorsal (D), dark grey; nasal (N), mid grey; ventral (V), light grey). Note that most respective tunings superimpose well, indicating that cone-tunings are approximately eye-region invariant. While significant regional variations existed in all cone-types (Discussion), these were generally very small when compared to across-cone-type differences. Accordingly, for further processing we used the averages of all zones of a given cone type. Note that heatmaps (d) are time-inverted to facilitate comparison to summary plots (f,g). Greyscale bars are in z-scores.*

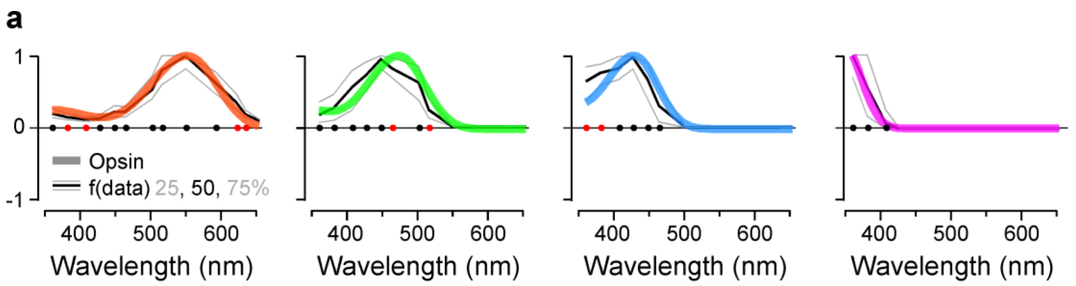


***Figure S2 | Statistical comparison of opsin tunings with in vivo cones-responses in absence of HCs (related to Figure 2). a**, exponential fits of the HC blockage to the opsin curves. Dots indicating for which wavelength the opsin curve lies within (black) or outside (red) 50% of the data distribution*

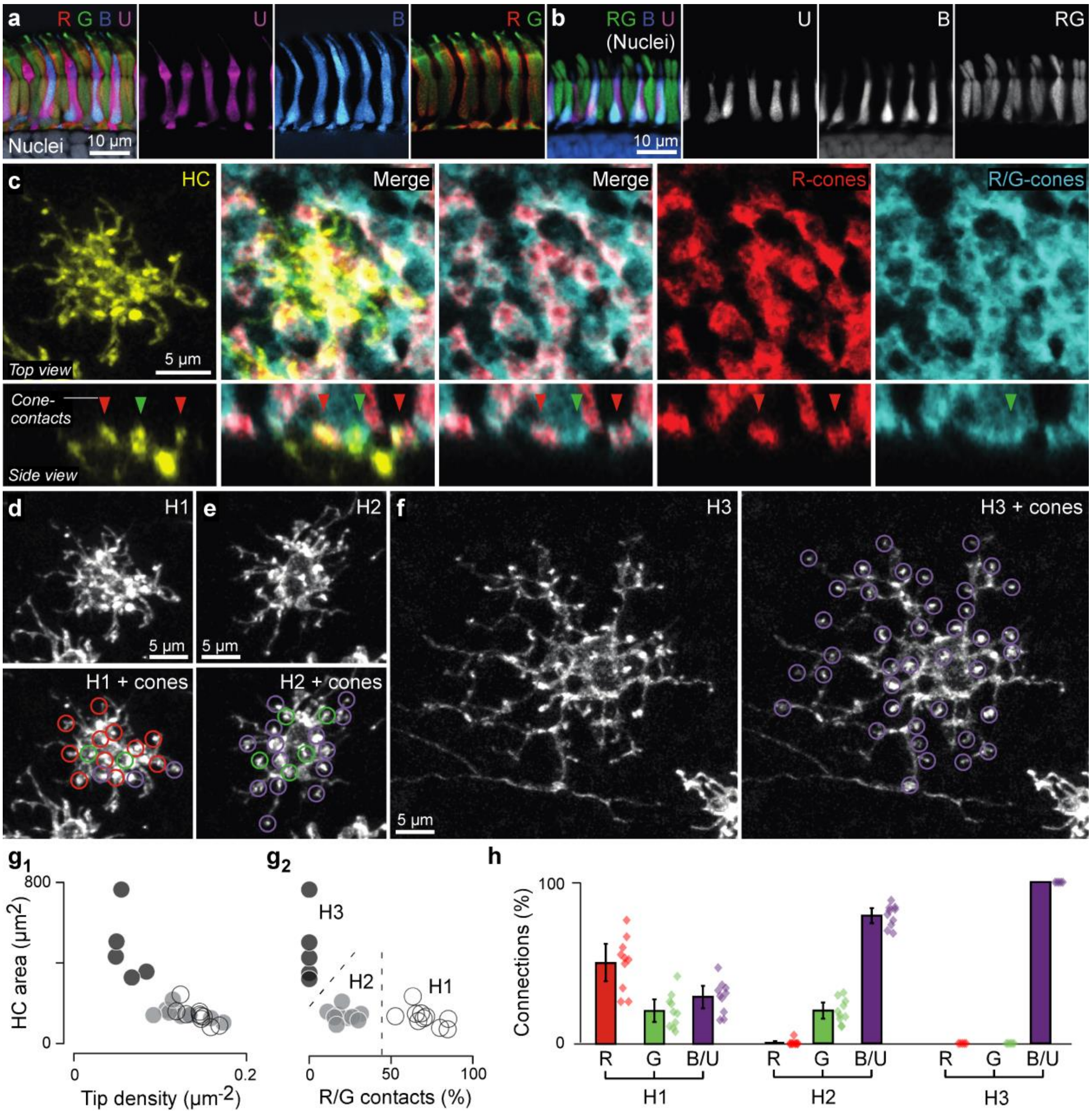


***Figure S3 | Cone-HC circuit connections by light microscopy (related to Figure 3). a,*** *vertical cross-section through outer retina, with the four cone-types individually labelled by the combination of transgenic labelling of cone types in Tg(Opn1sw1:GFP, Opn1sw2:mCherry, thrb:Tomato) and zpr-1 antibody immunostaining (red, green, blue, magenta, as indicated) on a background of a DAPI nuclear stain (grey) (Methods).* ***b****, as (a) but now DAPI signal in specific cone types were extracted.* ***c****, Example single HC randomly labelled by plasmid injection into one-cell stage eggs to express membrane targeting YFP, with cone pedicles of red cones (Tg(thrb:Tomato), red) and both red- and green cones labelled (zpr-1 antibody immunostaining, cyan). This allowed directly attributing each HC contact with red-, green-cones (e.g. arrowheads) or others (blue- or UV-cone).* ***d-f****, example single HCs identified as H1 (d), H2 (e) and H3 (f), with cone-type contacts indicated.* ***g****, HC dendritic area in relation to tip density ($g_1$) and percentage of red- and green-cone contacts ($g_2$) for n = 25 HCs allowed splitting HCs into 3 groups, here allocated to H1-3 as indicated.* ***h****, Relative cone contributions to the three HCs.*

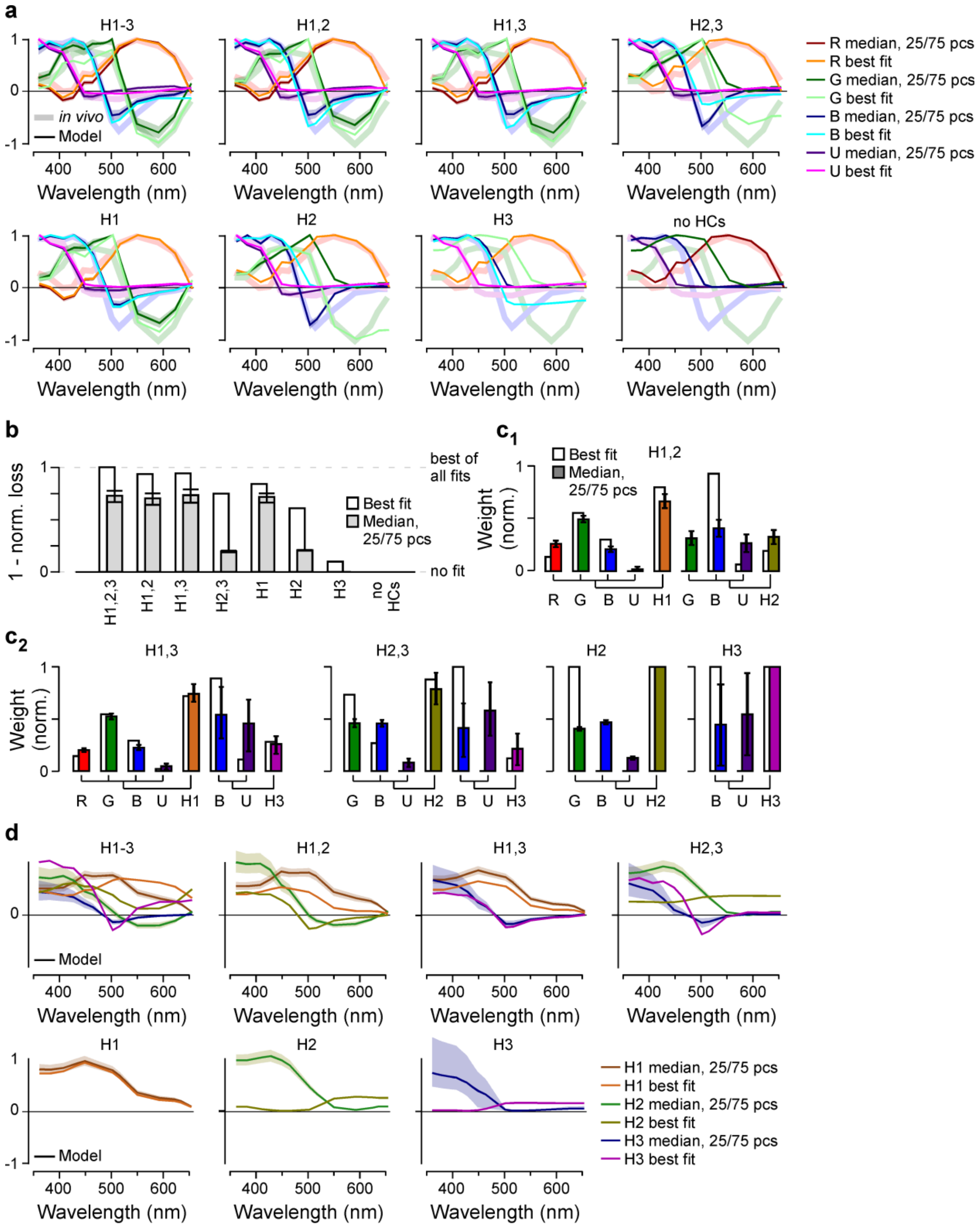


***Figure S4 | Additional HC model quantification (related to Figure 4). a**, cone tuning functions (as in Fig. 2b) that emerge from models comprised of different HC combinations as indicated, with best fit, median and 25/75 percentiles plotted on top of the measured in vivo cone tunings (thick shaded lines). **b-e**, normalised loss (b), distribution of weights (c) and emergent HC tunings (d) for all modelled HC-combinations (as Fig. 4c-e, respectively).*

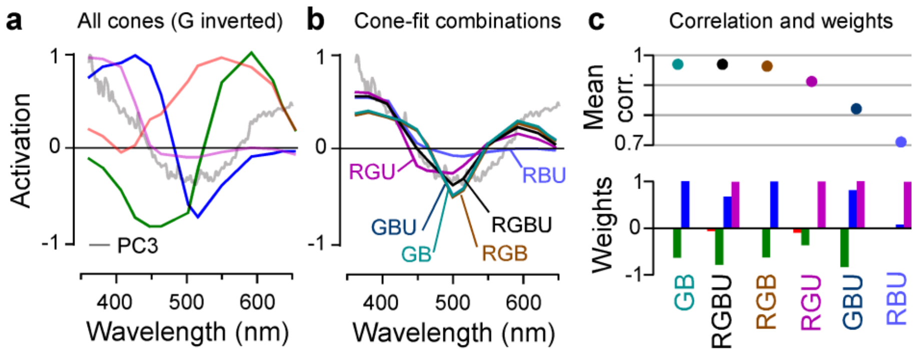


***Figure S5 | Linking zebrafish cone-tunings to PC3 (related to Figure 5). a**, mean in vivo cone-spectra superimposed on PC3 (from Fig. 5i, n), with green-cone tuning y-inverted for illustration. **b,c**, linear combinations of cone-tunings as indicated fitted to match PC3 based on least squares (b), and comparison of their performance in doing so (c). Note that GB, RGBU and RGB combinations all yield similar quality fits (ρ~0.97, top). In each case green-cones needed to be inverted relative to blue-cones (weights, bottom). Validation is based on the test data, as in Fig. 6a,b.*

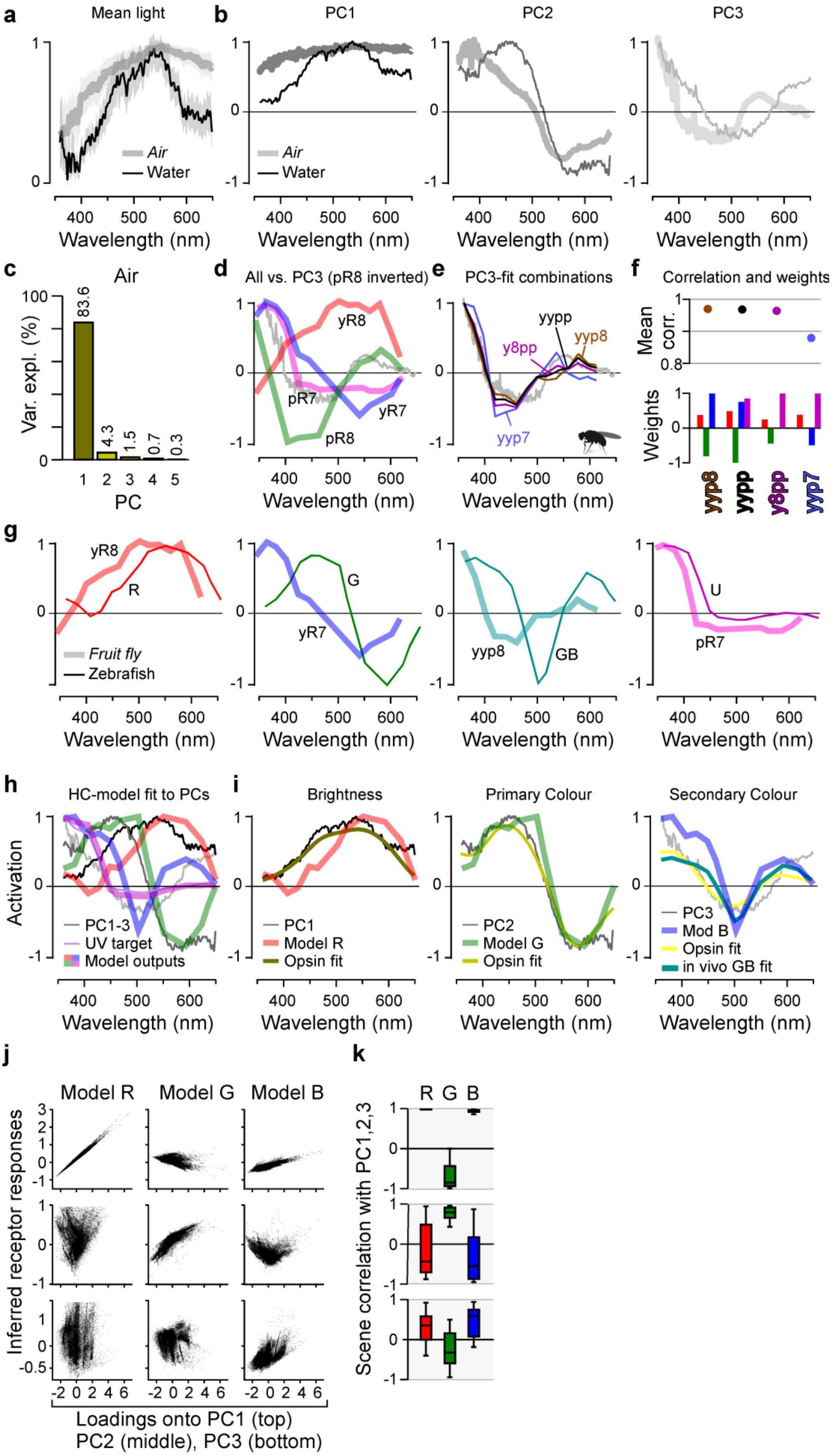
a
Mean light
Air
Water
Wavelength (nm)
b
PC1
PC2
PC3
c
Air
Var. expl. (%)
83.6
4.3
1.5
0.7
0.3
PC
d
All vs. PC3 (pR8 inverted)
yR8
pR7
yR7
pR8
e
PC3-fit combinations
yypp
y8pp
yyp8
yyp7
f
Correlation and weights
Mean corr.
Weights
yyp8
yypp
y8pp
yyp7
g
yR8
R
G
yR7
yyp8
GB
U
pR7
Fruit fly
Zebrafish
h
HC-model fit to PCs
Activation
PC1-3
UV target
Model outputs
i
Brightness
PC1
Model R
Opsin fit
Primary Colour
PC2
Model G
Opsin fit
Secondary Colour
PC3
Mod B
Opsin fit
in vivo GB fit
j
Model R
Model G
Model B
Inferred receptor responses
Loadings onto PC1 (top)
PC2 (middle), PC3 (bottom)
k
R G B
Scene correlation with PC1,2,3

***Figure S6 | Spectral processing in Drosophila, and a linear HC network fails to produce PC-like tuning in zebrafish blue-cones (related to Figure 6). a****, Mean±SD of the full spectrum of natural light in air (grey) and water (black) and* ***b****, the respective first three principal components that emerge. Note that “air-PCs” are systematically short-wavelength shifted compared to “water-PCs”.* ***c****, Variance explained by the first five PCs in the air-dataset.* ***d-f****, (as Fig. S52), here for fly R7/8 photoreceptors fitting to “air-PC3”. Note that the best fits opposed yR7+yR8 against pR8 (hence “yyp8”). Validation on the test data, as in Fig. 6a,b.* ***g****, superposition of “functionally homologous” fly- and zebrafish-photoreceptor tunings that capture PC1, PC2, PC3, respectively (from left to right). The “unused” zebrafish UV-cones and fly pR7 are superimposed in the final panel. Note that these four Drosophila curves are reminiscent of relatively short-wavelength shifted versions of the zebrafish curves.* ***h,*** *Horizontal cell model fit (cf. Fig. 4) when optimised to match red-, green- and blue-cones to PC1, PC2 and PC3, respectively. UV-cones were fitted to their own in vivo spectrum (as before), and* ***i****, respective matches superimposed. As before (cf. Fig. 5n), log-opsin-fit curves are added for reference.* ***j****, Comparison of measured in vivo tunings (thin lines) with respective HC-model outputs (thick lines).* ***k****, Evaluation of the above HC-model output tunings of red-, green- and blue-cones for capturing PCs1-3 (cf. Fig. 6a,b). Note that blue-cones still correlate with PC1 rather than PC3, while now green-cones fail to capture PC2.*

| | H1 weights (%) | | | | | H2 weights (%) | | | | H3 weights (%) | | |
|---|---|---|---|---|---|---|---|---|---|---|---|---|
| | R | G | B | U | H1 | G | B | U | H2 | B | U | H3 |
| D | 17.8 | 61.1 | 21.1 | 0 | 52.5 | 52.3 | 47.7 | 0 | 32.8 | 83.3 | 16.7 | 14.7 |
| N | 26.5 | 47.2 | 21 | 5.3 | 48.6 | 68.6 | 31.4 | 0 | 26.8 | 100 | 0 | 24.7 |
| AZ | 43.3 | 31.2 | 25.5 | 0 | 38.4 | 71.1 | 26.7 | 2.2 | 26.8 | 100 | 0 | 34.8 |
| V | 27.9 | 46.2 | 19.3 | 6.6 | 43.6 | 50.4 | 49.6 | 0 | 31.7 | 72.6 | 27.4 | 24.7 |
| All | 25.7 | 49.7 | 19.4 | 5.2 | 55.8 | 64.9 | 35.1 | 0 | 33.5 | 90.3 | 9.7 | 10.7 |

***Supplemental Table S1 – related to Figure 7.*** *Horizontal cell model outputs when individually computing spectral matches based on recordings taken only from one eye-region at a time. We used region specific cone recordings (Fig. S1g) as target for the model output as well as region specific model input from the experimental HC blocked condition (Fig. 2b). The relative weights of the best fits are shown, “all” refers to the data presented in Fig. 4d, “best fit” as in Fig. 4d, the posteriors (not shown) allow for some variability of the model weights, but the overall connectivity motifs stay the same across eye-regions.*
